# Lineage commitment pathways epigenetically oppose oncogenic Gαq/11-YAP1 signaling in dormant disseminated uveal melanoma

**DOI:** 10.1101/2024.03.05.583565

**Authors:** Rama Kadamb, Melisa Lopez Anton, Timothy J. Purwin, Lornella Seeneevassen, Vivian Chua, Francis Waltrich, Jessica L.F. Teh, M. Angela Nieto, Takami Sato, Mizue Terai, Sergio Roman Roman, Leanne De Koning, Deyou Zheng, Andrew E Aplin, Julio A Aguirre-Ghiso

## Abstract

Uveal melanoma (UM) can remain in clinical dormancy for decades only to later produce lethal metastases. Using Gαq/11^mut^/BAP1^wt^ UM xenograft models and human metastatic samples, we identified NR2F1 as a key inducer of UM disseminated cancer cell (DCC) dormancy. Dormant UM DCCs upregulate NR2F1, neural crest genes and, along with suppression of proliferation programs, NR2F1 silences YAP1/TEAD1 transcription by altering histone H3 activation marks. YAP1 can reciprocally repress NR2F1, but inhibiting Gαq/11 signaling or activating NR2F1 can arrest UM growth. NR2F1 knockout led to dormant DCC awakening and liver metastatic growth. NR2F1 and YAP1 inverse expression was confirmed in human livers carrying UM solitary, small DCC clusters as well as large metastases. Intriguingly, RNA-seq and Cut&Run analysis revealed that NR2F1 short-circuits oncogene signaling by repressing multiple G-protein signaling components. Our work provides previously unrecognized mechanistic insight into UM DCC dormancy and potential pathways for interception.

**Statement of significance:** NR2F1 epigenetically suppresses genes associated with G-protein signaling, cell cycle, and YAP1/TEAD1 pathways, inducing dormancy in uveal melanoma (UM) disseminated cancer cells. This study unveils novel markers for UM dormancy and reactivation, positioning NR2F1 as a promising target for intercepting residual and UM metastatic disease.

## Introduction

At the time of diagnosis, less than 5% of uveal melanoma (UM) patients have evidence of metastasis. However, 50% or more of UM patients will ultimately succumb to advanced metastatic disease ^1,2^. The liver is the predominant organ of metastasis ^3,4^. Standard chemotherapies rarely induce clinical responses in patients with macro-metastasis ^5–7^ and patient 1-year survival is 30-50% ^8,9^. A few drugs (e.g., tebentafusp and melphalan) have been shown to provide benefit in metastatic UM ^10,11^ but they only increase overall survival by few months. Thus, preventing the occurrence of metastatic disease remains an urgent unmet need for effective therapeutic strategies for UM.

Activating mutations (typically Q209) in alpha subunits of the heterotrimeric G proteins, *GNAQ* and *GNA11* were the first described driver mutations in UM and are found in 80-94% of UM ^14–17^. Risk of metastatic disease and UM associated survival is strongly correlated with mutation in key additional genes ^12,13^. Mutation of either *SF3B1* or *EIF1AX*, are associated with a favorable prognosis ^17,18^. Monosomy of chromosome 3 is associated with poor prognosis ^19^. Inactivating mutations of *BAP1* (<u>B</u>RCA1-<u>a</u>ssociated <u>p</u>rotein 1) on 3p21.1 are found in 32-50% of primary uveal melanomas ^20–22^ and associate with a higher likelihood of metastasis ^23,24^. *BAP1* encodes a de-ubiquitinating enzyme that removes mono-ubiquitin from histone H2A leading to activation of gene transcription ^25,26^. BAP1 is also implicated in DNA repair, cell cycle progression and cell differentiation ^27–29^. Furthermore, high expression of PRAME (preferentially expressed antigen in melanoma) is associated with metastasis ^30^. It is also evident from clonal phylogenetic analysis and autopsy studies that in a significant proportion of patients, UM can disseminate very early in evolution ^31–33^ and colonize multiple sites, with the liver being the site with highest frequency of eventual metastatic outgrowth.

A long-standing question in UM is the mechanisms that allows UM cancer cells to display a remarkable delayed time to metastasis, a phenomenon already recognized before the early 60’s with case reports dating back to the 40s’and earlier ^34^, but the mechanisms have remained unknown. Metastases often take years-to-decades to grow out and do so without recurrence at the primary site, highlighting early dissemination and tumor cell dormancy at distant sites as reported for UM and other cancers ^33,35–37^. How UM DCCs persist in this dormant form for so long, particularly in the liver, is a key clinical problem. UM liver dormancy is supported by studies in which fluorescently labeled cells were tail vein-injected into mice and non-proliferative single cells were identified in the liver ^38^. However, the mechanisms underlying delayed relapse and eventual regrowth remained unknown.

Melanocytes in the eye are derived from neural crest cells ^39,40^. We identified lineage commitment transcription factors (TFs) such as DEC2/BHLHE41 in the muscle and COUP-TFI/NR2F1 in neural crest cells, and primed pluripotency regulators like ZFP281 in the blastocyst as key regulators of dormancy in various cancer models ^41–43^. Interestingly, NR2F1 is specifically involved in lineage commitment of neural crest cells and differentiation during eye development ^44,45^. The above-mentioned TFs are regulated by upstream autocrine or niche-derived signals such as retinoic acid, TGFβ2, BMP7 and WNT5A. Downstream TFs such as NR2F1 and ZFP281 induce quiescence by upregulating a genome wide repressive chromatin state and cyclin-dependent kinase (CDK) inhibitors (e.g., p27^Kip1^,p16) and, thus, blocking G1/S transcriptional programs ^42,46^. In cutaneous melanoma a dormant phenotype is associated with phenotype switching linked to activation of neural and mesenchymal programs that cause growth arrest. Further, therapy-induced residual “persister” disease in cutaneous melanoma was also linked to adaptive transcriptomic changes and epigenetic remodeling ^47^. Whether similar mechanisms are also operational in UM dormancy was unknown until now.

Here we reveal, using xenotransplantation and unbiased isolation of quiescent human UM DCCs from the liver, that in Gαq/11^mut^/BAP1^wt^ human uveal melanoma cells, NR2F1 is a key regulator of dormancy of DCCs. Mechanism analysis revealed that dormant UM DCCs upregulate NR2F1 expression while silencing YAP1/TEAD1 transcription and growth programs. NR2F1 and YAP1 mutually antagonize their expression and NR2F1 binding to the YAP1 promoter that displays a loss of histone H3 activation marks and thus short-circuits oncogene signaling. The inverse correlation between NR2F1 and YAP1 was confirmed in human livers carrying solitary and small DCC clusters as well as large metastases. Importantly, agonist mediated NR2F1 activation or targeted inhibition of Gαq/11^mut^ enhances NR2F1 expression, blocking UM growth. UM solitary DCCs in mice and humans upregulated NR2F1 and *in vivo* CRISPR KO of NR2F1 results in dormant DCC awakening and massive liver metastatic disease. Our research provides explanation to the long-held mystery of UM dormancy, and it highlights the way through which dormancy pathways can be strong enforcers of lineage commitment and partial differentiation growth arrest programs that can epigenetically override apparently irreversible oncogenic programs caused by driver mutations.

## Results

### Long-lived dormancy of human UM DCCs in mouse livers

Clinical experience suggests that disseminated UM cells persist in livers for extended periods and, even as metastatic lesions, exhibit slow proliferation ^30,48,49^. To model this, we engineered human Gαq/11^mut^/BAP1^wt^ OMM1.3 UM cells to dually express H2B-GFP from an inducible TET-ON promoter and Td-Tomato from a constitutive promoter, creating the OMM1.3-QR line (OMM1.3 <u>q</u>uiescence reporter) (Fig. 1A). The H2B-GFP system stably labels nucleosomes in quiescent cells for long periods as reported for hematopoietic stem cells ^50^. Proliferating cells dilute the H2B-GFP signal, becoming H2B-GFP^-^/Td-Tomato^+^ (QR^-^), while non-proliferative cells do not turn over their nucleosomes and retain the signal as H2B-GFP^+^/Td-Tomato^+^ (QR^+^) (Fig. 1A). In 3D organoid cultures, Dox-induced OMM1.3-QR cells seeded as solitary cells cycled slowly, with ∼25% and ∼18% of label-retaining cells persisting at 7- and 14-days post-seeding, respectively (Fig. S1A). These results demonstrate that the H2B-GFP label retention system in OMM1.3 cells can identify slow-cycling/dormant-like cancer cells *in-vitro*.

**Fig. 1.**
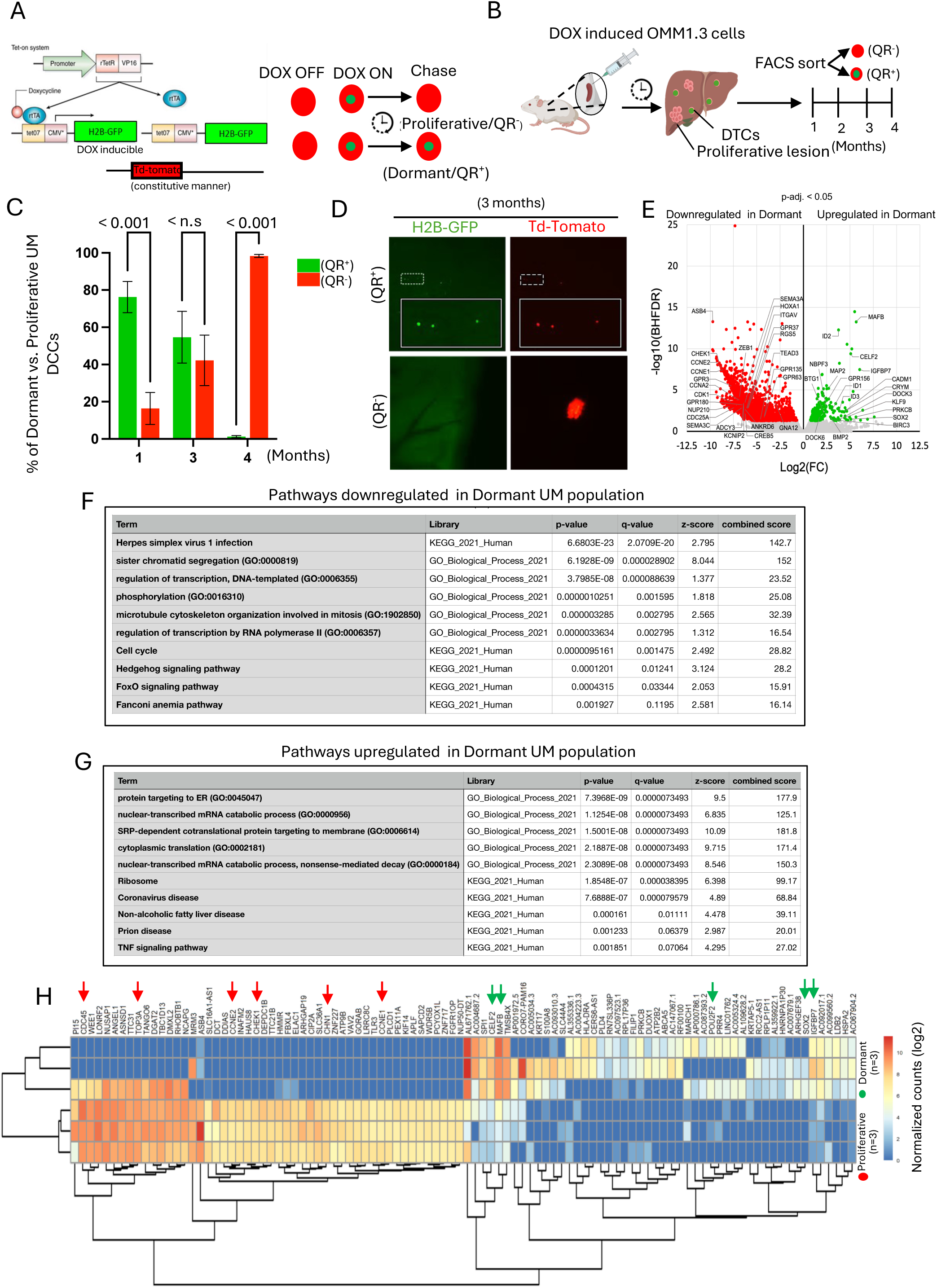
Long-lived dormancy of human UM DCCs in mouse livers: A) Scheme depicting the reporter construct used to generate OMM1.3 quiescence reporter (QR) cell line. B) Scheme depicting the in-vivo experimental strategy for intra-splenic injection, liver colonization and timeline for isolating QR^+^/QR^-^ cells. C) Percentage of dormant (QR^+^) and proliferative (QR^-^) UM cells disseminated to the liver determined by FACS sorting p-values calculated using multiple t-test. D) Representative images of the liver surface showing solitary OMM1.3-QR^+^ DCCs (Top) or OMM1.3-QR-proliferative micro-metastasis (Bottom) after 3 months intrasplenic injection. E) A volcano plot showing differentially expressed genes in dormant OMM1.3-QR^+^ and proliferative OMM1.3-QR^-^ sorted cells at 3-month time point (N=3). F&G) Pathways upregulated and downregulated in dormant OMM1.3-QR^+^ DCCs using Enrichr GO and KEGG term (cut off log2FC= +/-1). H) Heatmap showing top 50 significantly upregulated and downregulated genes in dormant and proliferative OMM1.3-QR cells (p-adj < 0.05). Arrows indicate cell cycle and transcriptional regulatory genes downregulated (Red) and neural-related genes upregulated (Green) in dormant OMM1.3-QR^+^ DCCs.

To test this tool *in vivo,* doxycycline-pulsed OMM1.3-QR cells were injected into spleens of nude mice for efficient spontaneous liver seeding ^51^. Animals were monitored for disease progression at 1, 2, 3, and 4 months (Fig. 1B). FACS analysis revealed that at 1 month, ∼80% of OMM1.3-QR cells were dual-labeled (OMM1.3-QR^+^) indicating quiescence. At 3 months the liver still contained ∼55% OMM1.3-QR^+^ quiescent cells and ∼45% OMM1.3-QR^-^ proliferated cells. By 4 months, ∼5% of cells were quiescent, while > 95% were proliferative (Fig. 1C, Fig. S1B & S1C). Epifluorescence microscopy at 3 months showed both solitary QR^+^ DCCs and QR^-^ micro-metastases, in extravascular spaces against green liver autofluorescence (Fig. 1D). Inspection of spleens at this time point revealed no visible UM growth but occasional solitary DCCs remained (data not shown). These findings indicate that upon liver arrival, the dominant phenotype of human UM DCCs is a slow- or non-proliferative state. Moreover, data at 3 and 4 months (equivalent to 9–12 years in humans) suggest that dormant cells coexist with proliferative lesions in the liver over prolonged periods.

### Differential regulation of neural and cell cycle programs in quiescent versus proliferative UM DCCs

OMM1.3-QR^+^ and QR^-^ cells were isolated from mouse livers via FACS at 1, 2, 3 and 4 months, followed by RNA-seq. These time points were chosen to maximize the detection of quiescent cells (Fig. S1C). RNA-seq data for QR^+^ cells at 1 and 2 months revealed very low level of human cells and the RNA-seq data was unreliable, possibly due to low numbers of cells recovered from the livers and/or contamination with host cells despite careful sorting. However, at 3 months, sufficient QR^+^ and QR^-^ cells were recovered from three mice (31, 34 and 36) to provide interpretable abundant RNA-seq data (Fig. S1C). Proliferative OMM1.3 cells cultured *in-vitro* served as a continuous proliferative control (Control group in Fig. S1B). Unbiased clustering by the number of expressed genes revealed that all three QR^+^ samples clustered separately from QR^-^ samples and closer to proliferative *in-vitro* OMM1.3 cells, supporting the notion that metastatic reactivation programs match the basic UM proliferative pathways (Fig. S2A). Further, differential gene expression (DEG) analysis from RNA-seq revealed that the majority of DEGs in OMM1.3-QR^+^ cells were downregulated (3045 genes, ∼90% of DEGs), suggesting a strong decrease in transcription of genes in various cellular processes during the 3-month dormancy (Fig. 1E, Fig. S2B). The genes associated with downregulated pathways showed significant reduction in the transcription of key cell cycle G1/S and G2/M transition genes confirming a growth arrest program in these cells (Fig. S1C).

Pathway analysis of downregulated genes in QR^+^ cells, using Enrichr database GO and KEGG terms, revealed significant suppression of transcriptional activation, cell cycle regulation, RNA Pol II activity, mitotic spindle organization and sister chromatid segregation (Fig. 1F, Table S1). Conversely, upregulated genes were associated with mRNA catabolism, activation of TNF signaling and protein targeting to the ER, consistent with ER stress pathways reported in dormant cancer cells ^52,53^ (Fig. 1G, Table S1). Analyzing the top 100 upregulated and downregulated genes further highlighted regulation of G2/M transition and GTPase activity, alongside upregulation of transcriptional repression and differentiation pathways (Fig. S1D & S1E). Genes linked to neural fate regulation, neurotransmitter signaling, and neurological disorders were notably upregulated (highlighted in green arrows), and genes linked to cell cycle regulation (highlighted in red) were downregulated (Fig. 1H). Since uveal melanocytes are derived from neural crest cells, it was consistent to find that a neural crest cell signature was activated in dormant UM OMM1.3-QR^+^ cells (Fig. 2A). This led us to hypothesize that proliferative OMM1.3 cells are in a de- differentiated state and when entering dormancy as OMM1.3-QR^+^ cells and upregulate neural lineage commitment genes as they exit from cell cycle. Given the role of NR2F1 in regulation of both melanocytic lineage and in maintaining dormancy, we tested the expression of NR2F1 in OMM1.3-QR^+^ and OMM1.3- QR^-^ cells. We did not detect NR2F1 expression in our RNA-seq analysis, possibly due to low transcript abundance from the rare OMM1.3-QR^+^ cells. However, qRT-PCR from an independent experiment with newly sorted OMM1.3-QR^+^ and OMM1.3-QR^-^ UM cells at 3 months showed > 5-fold upregulation of NR2F1 compared to proliferative UM DCCs (Fig. 2B). This observation was further strengthened by *in vivo* data using high resolution confocal microscopy, where we found selective upregulation of NR2F1 in the nuclei of solitary UM DCCs and < 8-cell clusters (Ki67^NEGATIVE^ dormant cells) vs. > 8-cell clusters (Ki67^POSITIVE^/proliferating cells) in the liver of mice, that were injected in the spleen using UM004 (Fig. S3A) or OMM1.3 cells (Fig. 2C & Fig. S3B). *In vitro,* we also categorized < 8-cell clusters as non-proliferative/dormant and > 8-cell clusters as proliferative due to low Ki67 expression in < 8-cell cluster and higher Ki67 expression in > 8-cell clusters in OMM1.3 cells grown in 3D (Fig. S3C). Importantly, the Ki67^NEGATIVE^ dormant cells in <8-cell vs. > 8-cell clusters being consistently Ki67^POSITIVE^ /proliferating cells also holds true in human UM patient samples that show low staining of Ki67 in < 8-cell cluster and high staining of Ki67 in > 8-cell clusters (Fig. S3D). Thus, solitary DCCs or clusters that only underwent up to 3 cell divisions can remain in a non-proliferative state in mouse models and human samples.

**Fig. 2.**
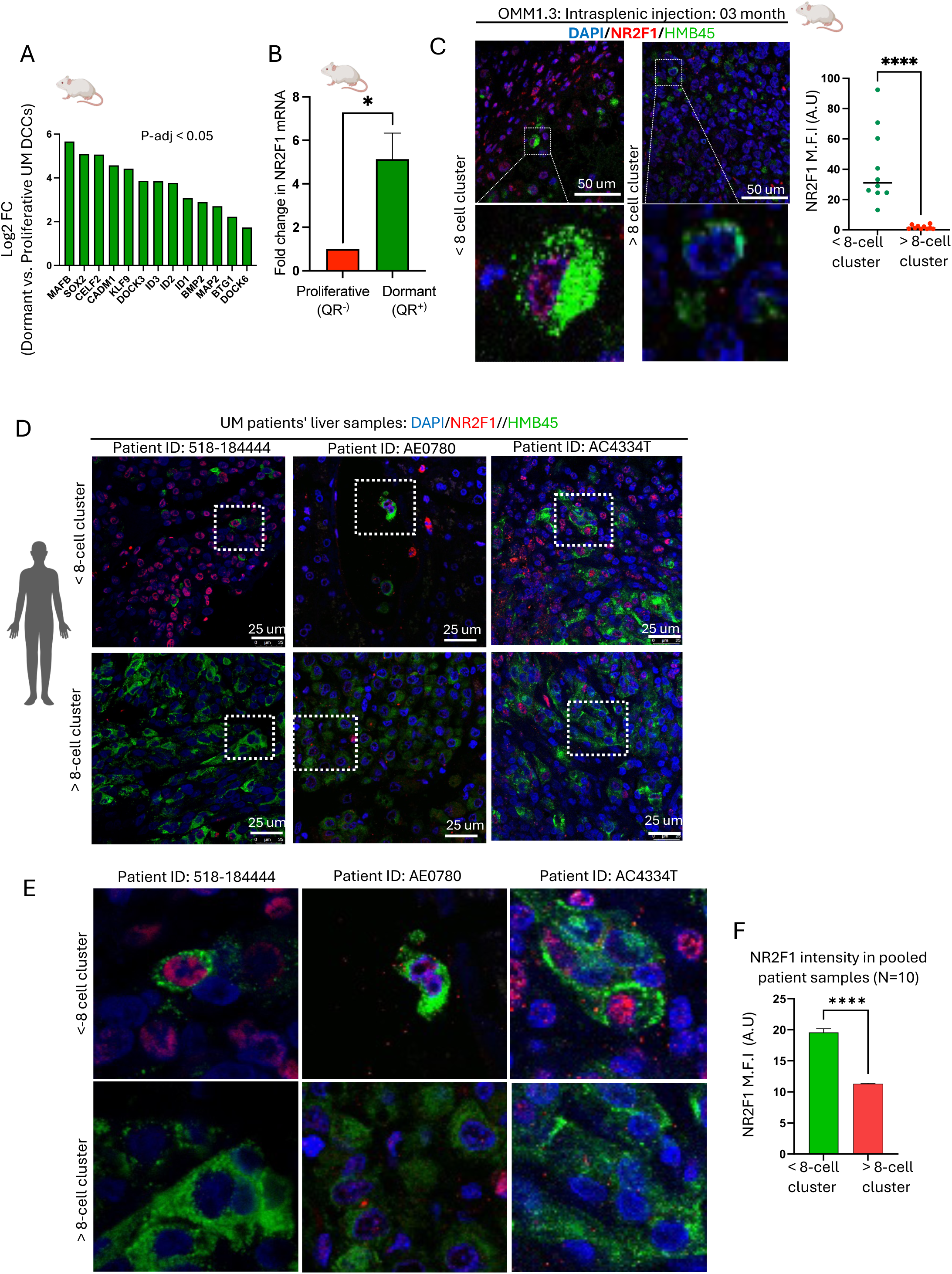
Dormant solitary UM DCCs express high NR2F1 and regulate neural and cell cycle programs: A) Expression of neural crest genes upregulated in OMM1.3-QR^+^ UM DCCs as revealed from RNA-seq DEG (p-adj. < 0.05). B) Fold change in NR2F1 mRNA expression in OMM1.3 QR^+^/QR^-^ FACS sorted cells isolated from the livers of mice after 3 months (student’s t-test; error bar= SD, n= 3). C) Representative IF images for expression of NR2F1 in xenografts of OMM1.3 cells in < and > 8-cell clusters disseminated to the liver after 3 months. HMB45 was used to track UM cells. Graph shows nuclear M.F.I (A.U) for NR2F1 in < and > 8-cell clusters. p-value calculated using Mann-Whitney test. D) Representative IF images for NR2F1 expression in < and > 8-cell clusters from three random metastatic UM patient samples. E) The dotted squares in panel D are shown as a zoom-in view. F) Quantification of nuclear M.F.I for NR2F1 in 10 independent UM patients’ sample and p-value is calculated using Mann-Whittney test.

RNA-seq, *in vivo*, and 3D model data consistently showed slow proliferation through downregulating cell cycle genes and low Ki67 staining. In addition, IF imaging showed that nuclei of slow proliferating OMM1.3 and UM004 cells in 3D that generate either solitary UM cells or <8-cell clusters express high NR2F1 and low Ki67 (Fig. S3E). In contrast, proliferative UM cell clusters of > 8-cells showed significantly lower frequency of NR2F1 expression. Importantly, NR2F1 expression was inversely correlated with Ki67 expression in both UM004 and OMM1.3 cells under *in vitro* and *in vivo* conditions (Fig. 2C, Fig. S3A & S3B and Fig. S3E). These observations imply that UM cells that enter quiescence as solitary DCCs and DCCs that undergo fewer than 3 divisions upregulate lineage commitment programs associated with neural crest identity and strongly downregulate proliferation programs that maintain dormant phenotypes

### Increased expression of NR2F1 in human UM solitary cells compared to proliferative/ metastatic masses co-existing in human livers

Liver samples from UM patients before overt metastases that would potentially carry dormant DCCs are unavailable. However, we have shown in mouse models, including Fig. 1 and 2, and patient samples that solitary or small cluster of DCCs can co-exist with larger metastatic lesions in the same organ and express dormancy markers ^54,55^. NR2F1 is key regulator of dormancy program in several cancer models ^42,56–58^ and we observed that dormant Ki67^NEGATIVE^ UM cells significantly enhance NR2F1 expression. This prompted us to test whether the association of increased NR2F1 in dormant DCCs also holds true in patients’ UM liver metastatic samples by quantifying NR2F1 expression in < 8-cell clusters (surrounding near the metastatic lesion or farther away in the normal liver tissue) and > 8-cell clusters. For this, patient samples (N=10) were used to co-detect NR2F1 and HMB45 expression, where the latter marker identifies UM cells in the HMB45^NEGATIVE^ background of liver cells. IF analysis showed that expression of NR2F1 is higher in solitary DCCs or < 8-cell clusters compared to > 8-cell clusters, which associated with low Ki67 levels (Fig. 2D, 2E & S3D). Interestingly we also found that hepatocytes in most patient samples also express NR2F1 (Fig. 2C & 2D). Overall, all analyzed patient samples showed that NR2F1 intensity is higher in solitary/isolated clusters compared to large clusters or the core of the metastatic lesions (Fig. 2F and Fig. S3F). In conclusion, NR2F1 is detected in UM samples and its upregulation correlates with the solitary and small cluster state. In contrast, proliferative lesions downregulated NR2F1 expression.

### YAP and NR2F1 gene expression is inversely regulated in dormant UM DCCs

In UM, Gαq/11 mutations are known to upregulate YAP/TAZ/TEADs pathway^59^ and a proliferation program in UM cells. We monitored YAP1 signaling genes in the quiescent OMM1.3-QR^+^ cells and OMM1.3-QR^-^ cells (as shown in Fig.1). Using DEG (p-value < 0.05) from RNA-seq data we found that *YAP1*, *TEAD1*, *TEAD3* and YAP1 pathway target genes (e.g., *NUAK2, DOC5*) were significantly downregulated in OMM1.3-QR^+^ vs -QR^-^ cells (Fig. 3A). Interestingly, genes related to the *Wnt* pathway (*GSK3B*, *WNT5A*) were also downregulated in the OMM1.3-QR^+^cells (Fig. 3A). The downregulation of YAP1 mRNA in OMM1.3-QR^+^ cells as compared to QR^-^ cells isolated from mouse livers after 3 months of growth was further confirmed by qRT-PCR (Fig. 3B). To confirm that dormant UM DCCs express low YAP1 and high NR2F1, we performed co-staining of NR2F1 and YAP1 in OMM1.3 and UM004 cells grown in 3D. IF analysis confirmed in both UM cell lines solitary or < 8-cell clusters expressed high NR2F1 and low levels of YAP1. For OMM1.3 cells, > 8-cell cluster expressed no or low NR2F1 but displayed very high nuclear YAP1 expression (Fig. 3C & 3D). UM004 cells also confirmed inverse relation to NR2F1 and YAP1 expression however, unlike OMM1.3 cells that showed strong nuclear YAP1 expression UM004 cells showed strong cytoplasmic and low nuclear YAP1 staining. Furthermore, among three UM cell lines tested, UM004 showed lowest expression of total YAP1 (Fig. S4A & Fig. S4B).

**Fig. 3.**
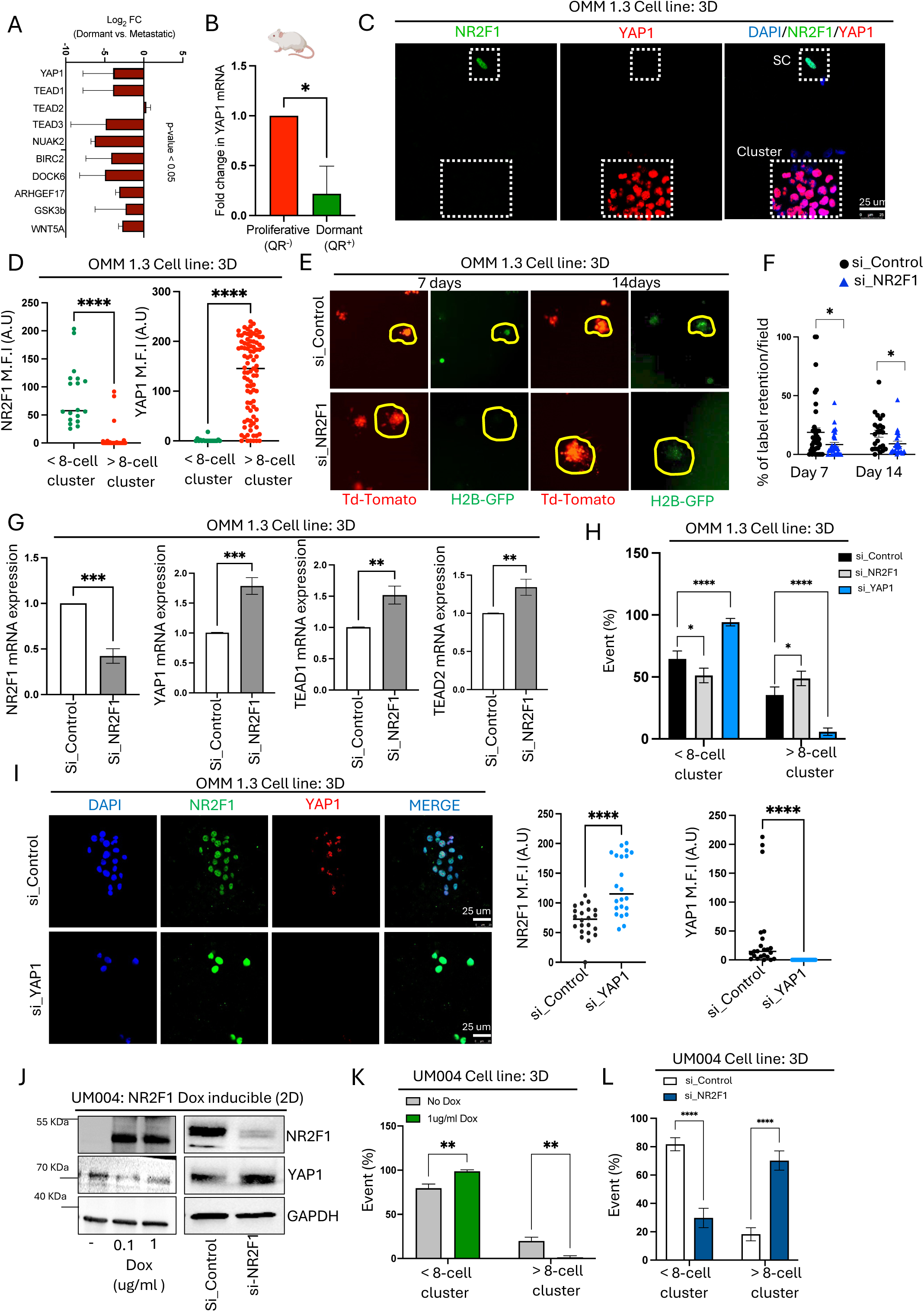
Antagonism between NR2F1 and YAP1 regulates proliferation of UM cells: A) Expression of YAP1, TEAD’s, and YAP1 downstream target genes in OMM1.3 QR^+^ cells as analyzed from RNA-seq DEG list (p-value < 0.05). B) Fold change in YAP1 mRNA expression in OMM1.3 QR^+^ (dormant) vs. QR^-^ (proliferative) cells at 3 months (student’s t-test, N=3, and * indicates p-value < 0.05). C) Representative IF images of OMM1.3 < and > 8-cell clusters co-stained for NR2F1 (green) and YAP1 (red) in 3D Matrigel. D) Quantification of nuclear NR2F1 and YAP1 M.F.I (A.U) in < and > 8-cell clusters formed by OMM1.3 cells in 3D and p-value calculated using Mann-Whitney test. E) Representative images of H2B-GFP and Td-Tomato expression in OMM1.3 QR colonies induced with doxycycline and transiently transfected with control or NR2F1-targeting siRNAs at 7- days and 14-days post-seeding. F) H2B-GFP label retention calculated at day 7 and 14 in OMM1.3-QR control and NR2F1 knockdown cells. p-values calculated using student’s t-test for each group. G) Relative mRNA expression of indicated genes in OMM1.3 control or NR2F1 knockdown cells. p-values calculated using student t-test. H) Graph showing % event of < and > 8- cell clusters formed by OMM1.3 cells upon transient knockdown of NR2F1 and YAP1. p-values calculated using one-way anova. I) Representative IF images (left) and graphs (right) showing expression and quantification of NR2F1 (green) and YAP1 (red) in colonies formed by OMM1.3 control and YAP1 knockdown cells at day 7. p-values calculated using Mann-Whitney test. J) Expression of NR2F1 and YAP1 in control (No Dox) and NR2F1 OE (Dox inducible) UM004 cells (left panel) and control and NR2F1 knockdown (siNR2F1) UM004 TR inducible cells respectively. K) Graph showing % event of < and > 8- cell cluster colonies formed by control and NR2F1 OE (dox inducible) UM004 cells after 7 days (n=4). L) Graph showing % events of colonies formed by control and NR2F1 KD UM004 cells at day 7 (n= 4). p-values were calculated using student’s t-test.

Loss of function assays in OMM1.3 cells using RNA interference (RNAi) targeting of NR2F1 resulted in accelerated growth of OMM1.3 cells as revealed by a reduced ratio of OMM1.3-QR^+^ to QR^-^ cells (Fig. 3E-F). Furthermore, *YAP1*, *TEAD1* and *TEAD2* mRNA expression was upregulated along with their downstream target genes *ANKRD1*, *CTGF1* and *CYR61* upon knock down of NR2F1 while expression of *TAZ* (*WWTR1*) gene remains unchanged (Fig. 3G & Fig. S4C). Loss of NR2F1 function in UM004 cell line also showed upregulation of *YAP1* with no significant changes in *TEAD1* and *TEAD3* expression (Fig. S5A).

YAP1 promotes growth and tumor progression of UM cells while the above data indicate that NR2F1 could maintain a dormant phenotype. Supporting this notion, we found that transient knockdown of YAP1 resulted in profound reduction in proliferative cell clusters and in growth suppression of UM cells in 3D culture (Fig. 3H). By contrast, NR2F1 knockdown enhanced UM colony growth. Interestingly, RNAi targeting of YAP1, which resulted in decreased UM colony growth, led to an increase in NR2F1 protein levels (Fig. 3I & Fig. S5B). At the protein level, inducing NR2F1 expression downregulated YAP1 expression while knockdown of NR2F1 enhanced YAP1 expression (Fig. 3J). In concordance with YAP1 and NR2F1 antagonistic expression and function we found that inducible expression of NR2F1 in UM004 cells strongly suppressed the ability of solitary UM cancer cells to transition into proliferative cell clusters while suppression of NR2F1 allowed the UM cells to proliferate (Fig. 3K & 3L). Further, at least in culture, we found that constitutive presence of NR2F1 is required to maintain growth suppression as removal of doxycycline from culture medium de-represses the growth inhibitory signal and enabled single UM cells to proliferate into big clusters (Fig. S5C).

Using a third UM line, 92.1 (primary lesion-derived UM cell line), we further confirmed the antagonistic relationship between NR2F1 and YAP1. Loss of function of YAP1 increased NR2F1 expression and concomitant decrease in > 8-cell UM clusters (Fig. 4A & 4B). In contrast, gain of function of YAP1 using a doxycycline inducible YAP1 expression system, resulted in decreased NR2F1 expression and increased > 8-cell clusters (Fig. 4C & 4D).

**Figure 4:**
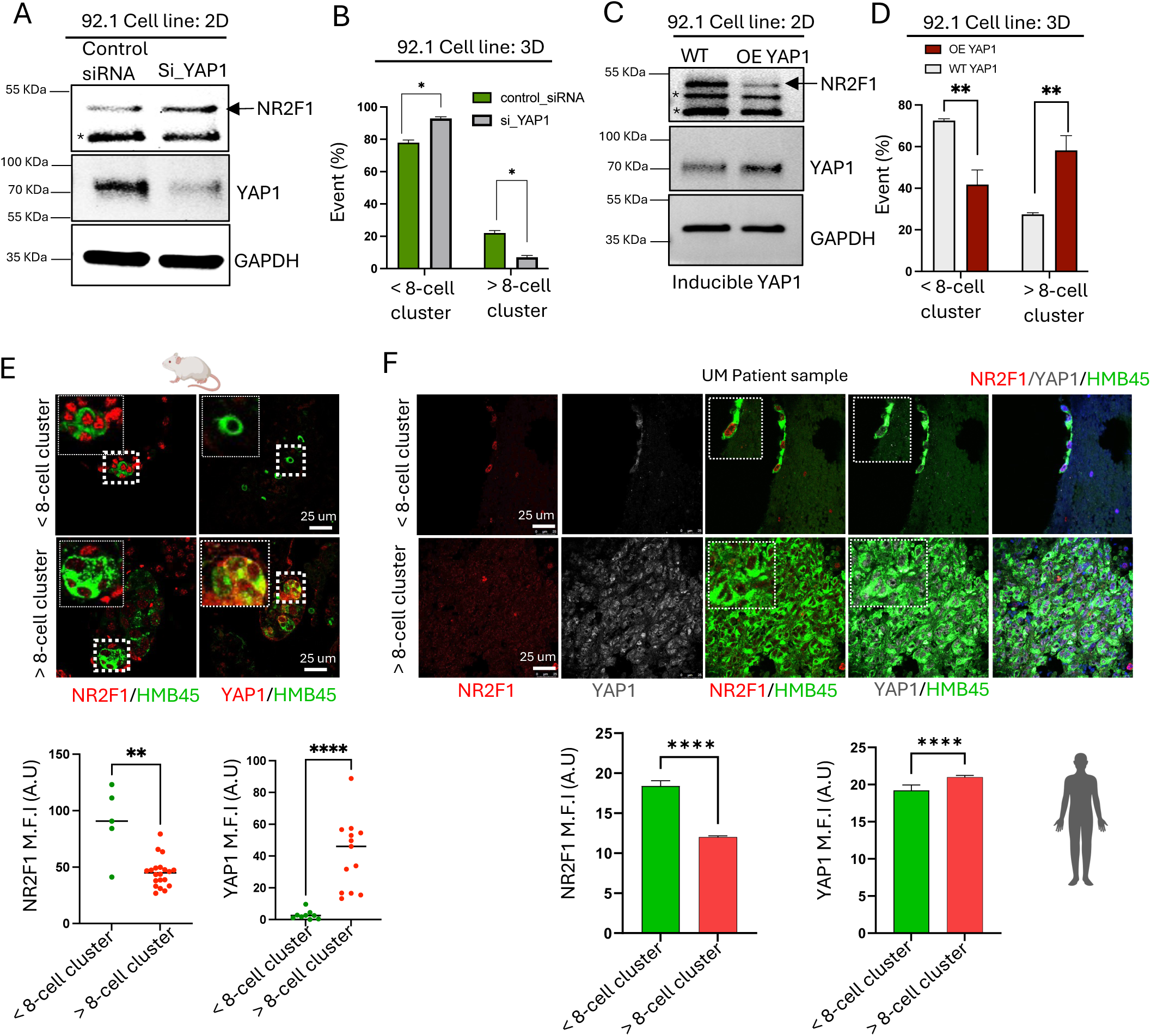
I**n**verse **expression of NR2F1 and YAP1 regulates UM proliferation in vivo and in human UM patients:** A) Expression of NR2F1 and YAP1 in 92.1 cells treated with either control or YAP1 siRNA. B) Graph representing % events formed by 92.1 cells treated with control or YAP1 targeting siRNA. C) Expression of NR2F1 and YAP1 in 92.1 cells treated with either control or NR2F1 siRNA. * Indicates non- specific band. D) Graph representing % events formed by 92.1 cells treated with control or NR2F1 targeting siRNA. p-values calculated using t-test. E) Representative IF images showing expression of NR2F1 and YAP1 3 months post UM004 cells dissemination to liver. Zoom-in images are shown in the same panel with marked boundaries. HMB45 was used to track UM cells. F) Representative IF images showing expression of YAP1 and NR2F1 in < and > 8-cell clusters in UM patient liver samples. Samples were co-stained for NR2F1 (red), YAP1 (grey) to show their expression in each UM cell. HMB45 was used to track UM cells. Graphs show the quantification of the nuclear M.F.I of YAP1 and NR2F1 (A.U.) in < and > 8- cell clusters in UM patient liver samples (N=5). p-values were calculated using Mann-Whitney test

To determine whether this apparent mutually antagonistic regulation between NR2F1 and YAP1 also existed in UM cell lines *in vivo* and in human UM patient samples, we stained liver sections recovered from mice injected in the spleen with UM004 cells and patient samples with liver metastases for co-detection of these two proteins. The data revealed a strong inverse correlation between NR2F1 and YAP1 in solitary/small, isolated clusters of UM DCCs and growing metastatic lesions in mouse liver (Fig. 4E). Antagonism between YAP1 and NR2F1 levels was confirmed in five UM liver metastasis patient samples by co-staining with NR2F1 and YAP1 antibodies in same section. Using independent staining in an additional five patient samples with NR2F1 and YAP1 antibody further confirmed an inverse relationship between NR2F1 and YAP1 expression profile in < 8-cell vs. > 8-cell clusters (Fig. 4F & Fig. S3F; S5D). There was significant inter-patient and intra-sample variation in YAP1 expression which reduced the delta between small and large clusters, but the difference was still significant. Nevertheless, our data support the notion that NR2F1 acts as an antagonist of the YAP pathway downstream of oncogenic Gαq/11 signaling and that it can suppress UM awakening and proliferation. However, the data also support that activation of YAP1 signaling can antagonize NR2F1 function.

### Assembly of YAP1-TEAD1 complex is negatively regulated by NR2F1

Our observations revealing that NR2F1 negatively regulates YAP1 led us to further explore its mechanism and significance. To this end we generated two independent NR2F1 CRISPR knock out (KO) OMM1.3 lines. Successful knockout of NR2F1 (NR2F1_KO1 & NR2F1_KO2) was confirmed by western blot (Fig. 5A). Specificity of knockout was shown by western blotting NR2F2, a paralog of NR2F1 that remains unchanged and only NR2F1 levels were affected by CRISPR mediated knockout (Fig. S6A). NR2F1 KO resulted in significant upregulation of YAP1, TEAD1 and CTGF1 along with mild upregulation of TAZ (Fig. 5A). In contrast, NR2F1 overexpression (OE) via an inducible system in UM004 cells led to downregulation of YAP1, CTGF1, TEAD1 and TAZ at a doxycycline concentration that we previously showed strongly inhibits UM cells proliferation (Fig. 5A & Fig. 3K). To our surprise, we observed that ERK1/2 phosphorylation (p-ERK1/2) was also increased upon loss of NR2F1 and reduced upon inducing NR2F1 expression, arguing that NR2F1 might negatively influence the Gαq/11 signaling also upstream of ERK1/2 and YAP1 through an unknown mechanism (Fig. 5B).

**Fig. 5.**
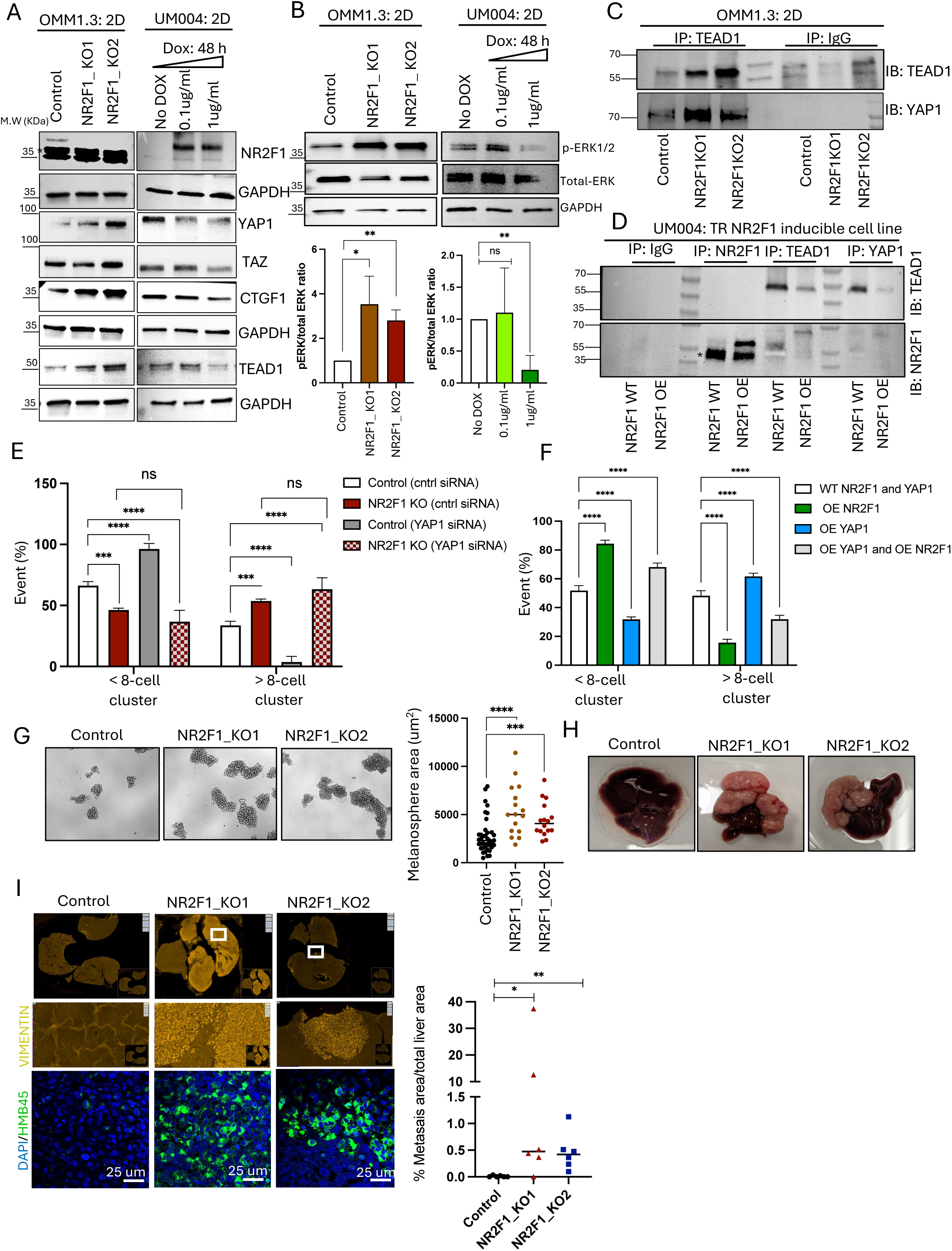
NR2F1 inhibits functional complex formation between YAP1-TEAD1 serving as a barrier to metastasis: A&B) Western blots showing expression of indicated proteins in control and NR2F1 knockout (KO) OMM1.3 cells, and UM004 control and dox inducible NR2F1 OE cells. * Indicates non-specific band. Lower panel in B represents densitometric quantification of ratio of pERK to total ERK in lysates from either OMM1.3 control and NR2F1 KO or UM004 control and NR2F1 OE cell lines. p-values calculated using student’s t-test. C&D) OMM1.3 control and NR2F1 KO cells and UM004 WT and NR2F1 OE cells as in panel A&B, processed for immunoprecipitation (IP) with the indicated antibodies followed by immunoblotting with anti- TEAD1, anti-YAP1 and anti-NR2F1 antibodies as indicated alongside each blot. E) Graph showing % event of < and > 8-cell clusters formed by OMM1.3 control and NR2F1 KO cells transiently transfected with YAP1 siRNA. p-values calculated using one-way anova and student’s t-test. F) Graph showing % event of < and > 8-cell clusters formed by UM004 WT, NR2F1 OE, YAP1 OE and both NR2F1 and YAP1 OE cells. p-values calculated using one-way anova. G) Representative images of melanospheres formed by control or NR2F1 KO OMM1.3 cells after 14 days. Right panel shows the quantification of average size of melanospheres in control and NR2F1 KO OMM1.3 cells. p-values calculated using Mann-Whitney test. H) Representative images of livers isolated 3 months after intra- splenic injection of OMM1.3 control and NR2F1-KO cells in nude mice. The images presented here represent the extreme case of liver metastatic burden, and not all animals showed such massive growth. I) Images showing vimentin expression in mouse livers injected with OMM1.3 cells after 3 months to quantify metastatic burden (upper row). The middle row is the zoom-in view of the marked area, showing the morphology of disseminated UM cells in the liver. The lower row shows IF images of the same sections stained with HMB45 (green) antibody and nucleus was stained with DAPI. The right panel shows the quantification of % metastatic area calculated using vimentin-positive cells. p-value calculated using Mann- Whitney test.

We next tested if the downregulation of YAP1 caused by NR2F1 was sufficient to reduce the assembly of the YAP1/TEAD1 complexes required for efficient transcriptional activity. Indeed, NR2F1 KO in OMM1.3 cells resulted in increased interaction between YAP1 and its co-activator TEAD1 as determined by co-immunoprecipitation (Co-IP) analysis (Fig. 5C). This increased interaction is specific to YAP1-TEAD1 as we found that YAP1 interaction with LATS1 remains unaffected upon NR2F1 deletion (Fig. S6B). NR2F1 upregulation via an inducible system in UM004 cells reduced the expression of YAP1 and TEAD1 and, consequently their interaction, as shown by Co-IP (Fig. 5D). These results show that NR2F1-mediated downregulation of YAP1 and TEAD1 caused a reduced availability of transcriptional complexes formed by these two TFs. We also demonstrate that NR2F1 does not interact with YAP1 or TEAD1 eliminating this possibility as an inhibitory function of NR2F1 on the hippo pathway TFs (Fig. 5D).

To confirm whether NR2F1 enforces dormancy primarily through suppressing YAP1, we carried out gain and loss of function of YAP1 in NR2F1 KO and NR2F1 OE background. RNAi mediated knockdown of YAP1 in OMM1.3 control cells resulted in significant suppression of UM cells growth in 3D Matrigel. In contrast, YAP1 knockdown in NR2F1 KO cells inhibit growth (Fig. 5E & Fig. S6C). On the other hand, overexpression of YAP1 in NR2F1 overexpressing UM004 cells suppressed colony growth compared to WT control cells; however, maximum growth inhibition still occurred in NR2F1 overexpression cells alone. (Fig. 5F). This suggests that in presence of high NR2F1 levels, YAP1 overexpression alone cannot completely override the dormancy program induced by NR2F1.

Based on these data, we tested whether loss of NR2F1 would unleash an unrestricted UM growth program. To test this hypothesis, we grew WT-NR2F1 and NR2F1 KO OMM1.3 cells in mammosphere medium to determine their “stem-like” capacity and how it is influenced by NR2F1 loss. These results showed that the KO of NR2F1 stimulated the growth of large UM colonies (Fig. 5G). We next tested whether this ability for enhanced growth would be able to interrupt the long lasting (months) dormant state of OMM1.3 cells in the mouse livers (Fig. 1). To test this hypothesis, we injected WT-NR2F1 and NR2F1 KO OMM1.3 cells via intrasplenic injection into nude mice and 3 months later solitary DCCs, micro- and macro-metastasis were scored by visual inspection of the livers and via histological and immunofluorescence analysis. Our data show that WT-NR2F1 OMM1.3 cells were, for the most part, unable to form visible metastases. IF analysis using human specific vimentin or HMB45 (to track UM cells) showed the perivascular persistence of solitary or small clusters of UM DCCs at 3 months (Fig. 5H & 5I) in the sections inspected. In contrast, NR2F1 KO OMM1.3 cells were able to overcome all growth restrictions keeping them in a solitary or small cluster state and formed metastatic lesions that invaded large sections of the liver (Fig. 5H & 5I). Consistent observations were made in both NR2F1 KO OMM1.3 cell lines. To rule out whether NR2F1 KO affects dissemination of UM cells after intrasplenic injection we assessed the number of UM cells in both control and NR2F1 KO groups that reached the liver 36 hours post intrasplenic injection. We found that NR2F1 KO does not influence liver seeding as there is no significant difference in the number of cells that seeded in the liver in control vs. NR2F1 KO groups and as expected in this short time frame all cells were quiescent (H2B-GFP^+^) (Fig. S6D). Hence, we conclude that the massive metastatic burden observed in the livers of mice injected with in NR2F1 KO UM cells must be the result of re-activation of the dormant UM DCCs after they seed the liver. We further conclude that NR2F1 upregulation drives a dormancy program that impedes UM metastasis, in part, by eliminating YAP1/TEAD1 complexes.

### NR2F1 represses transcription of the YAP1 gene and drives chromatin remodeling changes to inhibit GPCR-linked gene expression and signaling

NR2F1 dependent changes in YAP1 expression suggested a transcriptional repression mechanism induced by NR2F1 that silenced YAP1 expression. In-silico analysis suggests YAP1 promoter has putative binding sites for NR2F1 indicating that it might regulate YAP1 gene activity. Using Chromatin precipitation (ChIP) assay, we found NR2F1 enrichment onto YAP1 promoter near the transcription start site (TSS) region (from -320bp to -1000bp from the TSS) in OMM1.3 UM cells (Fig 6A). We further confirmed using qPCR that NR2F1 enrichment in the YAP1 promoter (highlighted as -320 bp region) was enhanced in UM004 cells upon NR2F1 induction (Dox induced), compared to WT (non-induced) cells. This enrichment was accompanied by a reduction in the accumulation of active marks H3K4me3 and H3K27Ac in the same region with no change in the H3K9me3 repressive mark levels (Fig. 6B). Accordingly, upon NR2F1 KO, we found enrichment of H3K4me3, H3K27Ac activation marks but not H3K9me3 repressive mark on histone H3 at promoter sites on YAP1 (near the TSS region) in OMM1.3 cells (Fig. 6C). These results suggest that NR2F1 epigenetically regulates YAP1 transcription through chromatin induced changes associated with transcription activating histone post-translational modifications.

**Fig. 6.**
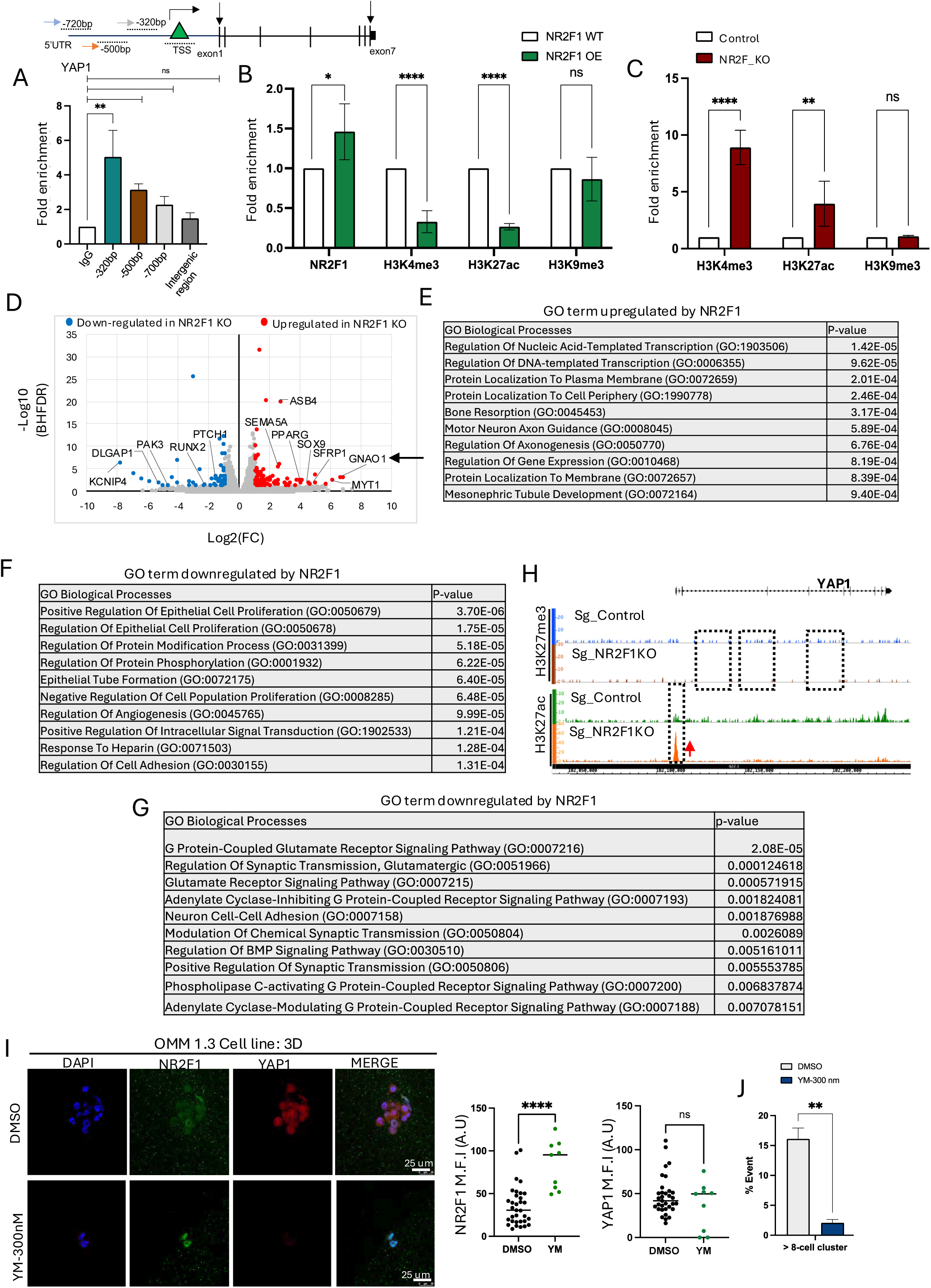
NR2F1 represses YAP1 gene transcription and drives chromatin remodeling changes to inhibit GPCR-linked gene expression and signaling: A) Schematic showing primer pairs used for scanning putative binding sites of NR2F1 on YAP1 genomic locus (1000 bp upstream of TSS). Lower panel shows fold enrichment of NR2F1 upstream of the TSS of the YAP1 promoter in OMM1.3 cells by ChIP-qPCR. Isotype-matched IgG was used as negative antibody control. B&C) ChIP-qPCR showing enrichment of histone carrying activation marks (H3K4me3 & H3K27ac) and repression (H3K9me3) marks at the YAP1 gene using primer pair (-320bp) in UM004 dox-inducible NR2F1 cells and OMM1.3 control and NR2F1 knockout cells, NR2F1 occupancy onto YAP1 genomic locus in UM004 cells was analyzed using primer pair (-320bp). p-values calculated using student’s t-test. D) Volcano plot showing DEGs in OMM1.3 control and NR2F1 KO cells cultured in 3D. E&F) GO term upregulated and downregulated as ranked by log2FC from top 100 upregulated and downregulated genes in NR2F1 KO OMM1.3 cells compared to control OMM1.3 cells using the Enrichr program. G) GO term repressed by NR2F1 based on downregulation of H3K27me3 peak in NR2F1 KO group compared to control (basal NR2F1) group. H) Distribution of H3K27me3 and H3K27ac peaks for YAP1 gene in control and NR2F1 KO condition (only one replicate is shown here). Tracks are normalized and scaled to the range by the integrative genomic viewer (IGV). I) Representative IF images showing expression of NR2F1 (green) and YAP1 (red) in colonies formed by OMM1.3 cells treated with 300nM YM for 14 days and corresponding quantification of NR2F1 and YAP1 MFI (A.U). p-value is calculated using Mann-Whittney test. J) Graph showing % event of > 8-cell clusters in OMM1.3 cells treated with DMSO and 300nM YM-254890.

Having established that NR2F1 represses YAP1 transcription, we conducted bulk RNA-seq to assess the NR2F1-regulated transcriptional program in OMM1.3 cells in 3D growth basal conditions. RNA-seq analysis was performed in NR2F1 WT and OMM1.3 KO cells cultured in 3D. We found significant changes in gene expression in control and NR2F1 KO cells (Fig. 6D &S7A). Analyzing the top 100 downregulated genes from the NR2F1 KO group by Enrichr GO terms showed that NR2F1 positively regulates genes involved in transcriptional regulation, motor neuron axon guidance, axonogenesis, ganglioside biosynthetic process and regulation of cytoskeleton organization (Fig. 6E, Table S2A). Downregulated genes in NR2F1 KO group included *RADIL*, *PLXNA3* and *SALL1*, that are associated with neural crest and axon guidance. Conversely, NR2F1 negatively regulated genes such as *NUP210* (linked to metastasis), *SFRP1*, and *PPARG* (associated with cancer progression), as reported by us and others and repressed biological processes involved in cell proliferation, angiogenesis, and protein modification (Fig. 6F, Table S2B).

Intriguingly, among the profoundly NR2F1-repressed genes was *GNAO1*, an activator of GPCR signaling known to interact with genes in the G protein signaling network required for proliferation (Fig. 6D). We further confirmed that GNAO1 transcript levels were upregulated in NR2F1 KO OMM1.3 cells while NR2F1 overexpressing UM004 cells suppressed GNAO1 mRNA levels (Fig. S7B & S7C).

Results from our ChIP and RNA-seq suggests that there appears to be a chromatin remodeling program under the regulation of NR2F1, aimed at interrupting Gαq/11 oncogenic signaling. To explore this hypothesis, we used Cleavage Under Targets and Release Using Nuclease (CUT&RUN) for analyzing enrichment of histone H3K27me3, H3K27ac and H3K4me3 modifications globally in NR2F1-WT or NR2F1-KO cells cultured in 3D assays for 7 days when the phenotypes become evident. The analysis did not find a large reorganization of chromatin modifications but revealed many chromatin regions with both increased (n=788) and decreased (n=635) H3K27me3 upon NR2F1 KO (Fig. S7D), with 57% of the changes located to promoters. Similarly, we also found 564 genomic regions with significant H3K27Ac change upon loss of NR2F1 (Fig. S7E), with 92% at promoter regions. Interestingly, the level of changes in H3K27Ac seem more pronounced than those in H3K27me3 (Fig. S7D & S7E); we were unable to capture significant peaks for H3K4me3 mark and therefore, this mark was not considered in the analysis.

As YAP1 is a downstream effector protein of GPCR signaling and we found YAP1 downregulation by NR2F1, we examined the distribution of H3K27me3 and H3K27Ac peaks onto YAP1. Analysis of peak distribution using IGV showed removal of H3K27me3 distribution at several positions while the intensity of H3K27ac peak was enhanced near the promoter region of YAP1 in NR2F1 KO cells (Fig. 6H). Gene ontology analysis from genes that showed loss of H3K27me3 peak in NR2F1 KO condition compared to control group revealed that NR2F1 represses G-protein glutamate receptor signaling and adenylate cyclase-inhibiting GPCR signaling (Fig. 6G, Table S3A).

Interestingly, NR2F1 KO repressed genes (that showed accumulation of H3K27me3 mark) showed that basal NR2F1 is involved in regulating cation transport and chemical synaptic signaling although its role in inducing a dormant DCC phenotype in UM is unknown (Fig. S8A, Table S3B). GO term analysis using genes that showed loss H3K27ac activation marks in NR2F1 KO revealed that basal NR2F1 upregulates biological processes related to nervous system development, differentiation, and the positive regulation of biosynthetic processes upon (Fig. S8B, Table S3C).

The data showing that YAP1 downregulation could result in NR2F1 upregulation and suppress colony proliferation led us to test whether direct inhibitors of the Gαq/11 activity (cyclic depsipeptide YM- 254890) could reduce UM proliferation and induce NR2F1. Treatment of OMM1.3 cells with 300 nM dose of YM-254890, increased NR2F1 expression with no significant decrease in YAP1 expression and suppressed UM growth (Fig. 6I & 6I). However, when each cell was analyzed in DMSO and YM group mean nuclear expression of YAP1 or NR2F1 showed antagonism (Fig. S9A) supporting that NR2F1 suppression is initiated by Gαq/11 activity. Further, the frequency of cells positive for NR2F1 increased in a dose dependent manner with YM-254890 treatment. In < 8-cell clusters, doses as low as 100 nM achieved almost maximal induction of NR2F1 in this population of cells. In contrast, > 8-cell clusters that presumably were already formed at the time the treatment was initiated, the effect was less pronounced but still highly significant with 300 nM dose showing the strongest induction of NR2F1 (Fig. S9B). This argues that both small and larger clusters respond to Gαq/11 inhibition to restore NR2F1 expression and dormancy onset. To further test the pharmacological approach to induce UM DCC dormancy, we treated OMM1.3 cells with C26, a novel NR2F1 agonist ^56^. C26 treatment increased expression of NR2F1 and reduced UM colony growth (Fig. S9C & S9D).

We conclude from our RNA-seq, ChIP-qPCR and CUT & RUN experiments that a multilayered program regulated by NR2F1 plays a major role in dormancy regulation and that regulators of G-protein signaling are being significantly repressed by NR2F1-induced chromatin changes. We further conclude that either inhibiting Gαq/11 activity and/or stimulating NR2F1 function might be alternatives to control UM metastatic disease.

## Discussion

The mechanisms behind the development of UM liver macro-metastasis, more than a decade after primary tumor treatment are still unclear ^62^. Dormancy of DCCs has been proposed to explain this long observed but poorly understood phenomena in metastatic UM. Our study aims to identify the mechanisms maintaining DCC dormancy, potentially leading to new metastatic UM treatments.

Using our H2B-GFP Tet ON/OFF label retention model for UM DCC dormancy, we successfully revealed that human Gαq/11^mut^/BAP1^wt^ UM cells can enter a prolonged dormancy state as solitary DCCs in the mouse liver equivalent to 9-12 years in humans. Previous studies revealed that Gαq/11 triggers the activation of the YAP1/TAZ pathway, which represents a major driver in UM progression ^59,63^. Analysis of transcriptional programs in OMM1.3-QR^+^ dormant and OMM1.3-QR^-^ UM DCCs also revealed that upon entry into dormancy the UM cells depart from a YAP1^hi^/TEAD1^hi^ state and acquire traits of neural lineage. Acquisition of the neural lineage is accompanied by silencing cell cycle progression, melanocytic and DNA synthesis programs. These changes are likely in response to a de-differentiation process from a melanocytic program towards a neural lineage, which is also positively regulated by NR2F1 during development ^44^.

YAP1 and TEADs have been characterized as targets of the Gαq/11 oncogenic signaling for UM growth ^63,64^. Thus, from a therapeutics point of view, it is an important finding that they are naturally downregulated during spontaneous dormancy of UM cells (OMM1.3-QR^+^ cells). Melanocytes that serve as precursor for nevi cells for UM initiation originates from neural crest cells ^65,66^. NR2F1 a nuclear receptor with only recently identified putative ligands (e.g., sphingolipids) ^67^, is a critical regulator of this lineage program and serve a pivotal role in cortical pattering during eye morphogenesis ^44,68,69^. Our data identified that among neural crest lineage genes, OMM1.3-QR^+^ dormant DCCs upregulate NR2F1 expression, previously linked to dormancy in breast, head and neck squamous, and prostate cancer DCC models ^58,70^. This allowed functionally determining that solitary DCCs or small DCC clusters (< 8-cells) that enter dormancy must acquire a YAP1^lo^/TEAD^lo^/NR2F1^hi^ program. Upregulation of NR2F1 could be detected in solitary or small clusters of OMM1.3-QR^+^ UM cells growing in 3D Matrigel, suggesting that the density of the UM cell cluster can make UM cells permissive for signals that turn on NR2F1 expression and a dormant phenotype. Basal NR2F1 expression in OMM1.3 cells appears to repress UM progression genes, facilitating dormancy entry in the liver and slow 3D proliferation. NR2F1 protein upregulation was observed in solitary UM cells, small clusters, and cells near the metastasis-liver boundary in both xenografts and human samples. These data were intriguing as they show that even advanced disease carries the ability to upregulate NR2F1 mostly when small clusters of UM cells or the edges of the metastatic lesions are in contact with the liver microenvironment. The metastasis core was mostly NR2F1 negative and inversely correlated with Ki67 expression. NR2F1 could potentially serve as a dormancy marker for UM DCCs or a therapeutic target for future reprogramming therapies^42,56–58^ .

The inverse expression of YAP1 and NR2F1 led us to explore how these co-activators/TFs might be interconnected at a regulatory level. First, we determined the importance of NR2F1 in enabling a dormant

phenotype as in 3D culture models, “melanosphere” cultures and in the livers *in vivo,* NR2F1 loss of function invariably led to enhanced growth. This was striking *in vivo* as loss of NR2F1 via KO led to relentless growth in the liver of cells delivered *via* intra-splenic injection or directly to the liver. Our data from three UM models indicates NR2F1 acts as a spontaneous barrier to metastatic progression by inducing dormancy, consistent with our previous findings in other cancer models showing NR2F1’s role in suppressing growth and metastasis burden^56,71^. High YAP1 activity reported to be critical for survival of UM cells ^64 59^. We found that invariably the YAP1 pathway was negatively regulated by NR2F1 and forced downregulation of NR2F1 led to a putative de-repression of YAP1/TEAD1 genes and activated YAP oncogenic signaling (Fig. 7). YAP1 expression was also absent in solitary cells in 3D models or in solitary UM DCCs *in vivo* and in humans or in small clusters but became detectable in metastatic cores in humans and large cluster in 3D cultures and in the mouse liver UM metastasis. Perhaps combined detection of YAP1 and NR2F1 levels in metastasis resected as a treatment may more firmly inform on disease progression and therapy responses in those patients. Surprisingly, the growth stimulation upon NR2F1 KO was independent of YAP1 at least in vitro, suggesting that other TFs and programs may take over the active growth after NR2F1 KO. It is possible, although we did not test it, that TEADs become major drivers of growth promotion independently of YAP1.

**Fig. 7:**
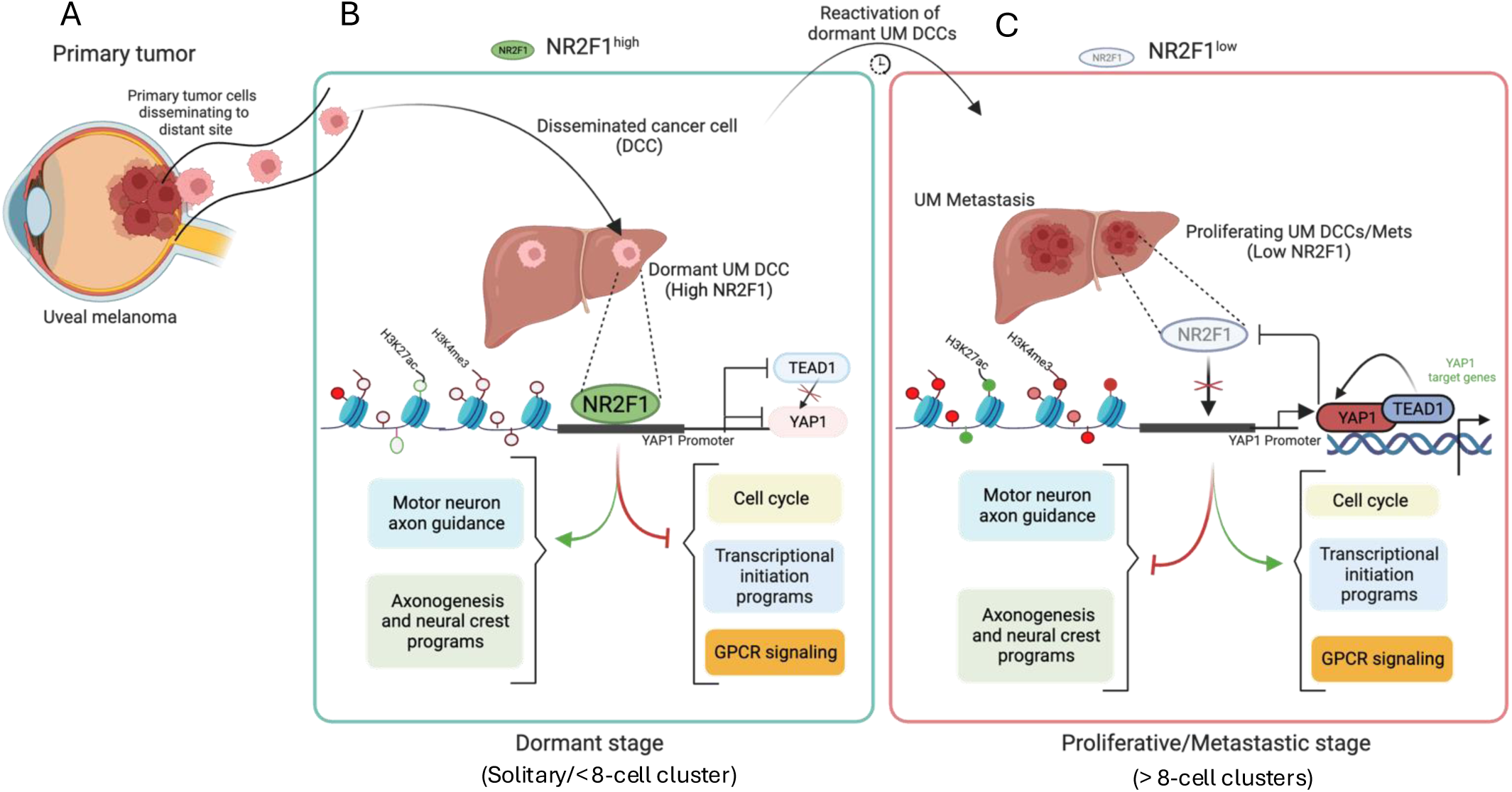
U**M cells disseminate to the liver and stay dormant for a prolonged period before manifesting as overt metastasis:** A) Dormant UM DCCs express high levels of NR2F1 which induces transcriptional repression of YAP1 by binding to its promoter B) Binding of NR2F1 inhibits recruitment of H3K4me3 and H3K27ac gene activation marks onto YAP1 promoter and globally represses programs related to cell cycle proliferation, transcriptional initiation and GPCR signaling while activating axonal guidance and neural crest migration pathways in dormant UM DCCs. C) With time, dormant UM DCCs lose NR2F1 expression and are reawakened to give rise to overt metastasis as NR2F1 binding onto the YAP promoter is reduced leading to activation of YAP1 oncogenic signaling. Also, YAP1 inhibits NR2F1 expression through unknown mechanisms causing activation of cell cycle proliferation, transcriptional initiation and GPCR pathways and inhibition of neuronal programs.

Our data revealed that NR2F1 can actively downregulate YAP1 and TEAD1 mRNA and protein expression. Detection of YAP1/TEAD1 complexes using co-immunoprecipitation showed that their downregulation by NR2F1 limits the availability of YAP1/TEAD1 transcriptional complexes required for the Gαq/11 oncogene growth promoting function (Fig. 7). We investigated whether inhibiting the Gαq/11 oncogene could reciprocally restore NR2F1 expression in UM cells. This was confirmed when we treated UM OMM1.3 cells with YM, a small molecule inhibitor of Gαq/11, which led a strong upregulation of NR2F1 followed by a strong growth arrest. As targeted therapies for UM and Gαq/11 oncogene develop, they may induce dormancy as a side effect. This may offer an opportunity for maintenance therapies using epigenetic therapies that maintain the NR2F1 program ^58^ and/or using NR2F1 agonists ^56^ that maintain its function during or following Gαq/11 inhibitor treatments.

We investigated how NR2F1 disrupts Gαq/11 oncogene signaling by restricting YAP1 and TEAD1 complexes. NR2F1, a nuclear receptor, has specific target genes and interacts with retinoic acid receptors to influence lineage commitment and differentiation.^72,73^. It has also been shown that NR2F1 can act as a transcriptional repressor ^74^. It has been shown that an entire cluster of transcriptional factors that regulates neural-crest and melanocytic differentiation bound YAP responsive elements ^75^. Our ChIP data identified that NR2F1 also binds to a specific region of the YAP1 promoter leading to a loss of histone H3 activation mark and reducing the rate of transcription of YAP1 (Fig. 7). NR2F1 didn’t alter histone H3 repressive marks, indicating additional signals may be needed for stable YAP1 repression.

RNA-seq data confirmed NR2F1 regulates genes involved in transcription, neural crest cell migration, axonogenesis, and biosynthetic pathways. Interestingly and supporting an anti-UM metastatic function of NR2F1 we found that genes linked to metastatic outgrowth such as *NUP210, WNT5A, SFRP1* and *PPARG* were repressed by NR2F1. This might explain the aggressive metastatic outgrowth observed upon NR2F1 downregulation and how NR2F1 may ensure dormancy of UM cells by limiting the response to these cues from autocrine or paracrine sources. Further, NR2F1 inhibits genes related to cell proliferation (*SEMA5A, SFRP1, WNT5A, IGF1, SOX9*), cell differentiation (*SFRP1, IGFBP3, PPARG, ASB4, SOX9, IGF1*) and epithelial to mesenchymal transcription (*WNT5A, SOX9, IGF1*) pathways. An unexpected, but in retrospective, logical result from our RNA-seq was that NR2F1 repressed a strong activator of GPCR signaling, *GNAO1* which is required for G-protein signaling for proliferation. GNAO1 is shown to promote growth in HCC and interacts with Phospholipase C (PLC) upstream of the MEK-ERK1/2 pathway activated in UM ^76,77^. In support of GNAO1 repression by NR2F1, we also found negative regulation of GPCR signaling by NR2F1 as revealed by the CUT & RUN analysis. NR2F1-led to the accumulation of histone H3 repressive marks in GPCR signaling genes associated with adenylate cyclase-inhibiting GPCR signaling and G protein-coupled glutamate receptor signaling pathways. Some of the GPCR pathway genes were already associated with metastasis promotion in different cancers (e.g., *GRM3, GRM8, RXFP1*) ^78–80^.

Our findings suggest a broader concept potentially applicable to various cancers: potent oncogenic signaling can be overridden by specific transcriptional programs. This supports the idea that in UM and other cancers, chromatin remodeling and transcriptional regulation (e.g., by NR2F1, ZFP281, or DEC2) can, under certain conditions and for extended periods, dominate over genetic driver mechanisms. Future research into a potential developmental link between NR2F1 transcriptional function and GPCR signaling activation could provide deeper insights into UM metastatic progression and reveal new therapeutic opportunities.

## Limitations of the study

Our human sample studies were limited by the lack of access to liver samples of UM patients in remission that would be ideal to detect *bona fide* dormant UM DCCs. We cannot determine whether the solitary UM cells or small clusters in the liver that were NR2F1^hi^/YAP1^lo^/Ki67^lo^ were cells that separated from the neighboring metastasis or arrived independently to the liver. While we cannot conclude these are truly early DCCs that would eventually find a metastatic growing lesion, the data support that NR2F1 along with YAP1 and Ki67 could be used to profile the presence of dormant DCCs in this disease perhaps after resection of single or oligo-metastasis or if liver samples were to be obtained during remission phases. We used a mouse model that only carries innate immune cells; thus, we cannot rule out the role of the adaptive immune system in controlling dormancy phases. Ongoing studies will reveal how innate immune cells regulate the pathways identified in this paper. The pathways associated with the genes linked to active histone modifications in NR2F1^hi^ UM cells revealed also a synaptic signaling pathway but how they may be linked to the induction of a dormant DCC phenotype is unknown. Future, studies are needed to test the therapeutic potential of inhibiting Gαq/11 activity or activating NR2F1 *in vivo* as well as inhibiting GNAO1 function and or activity.

## Materials and methods

### 1. Cell culture, transfection, and drug treatment

The human OMM1.3 and UM004 (genetically engineered to knockout or overexpress NR2F1 upon Doxycycline treatment) cell lines were maintained in DMEM-F12, and 92.1 cells were maintained in RPMI supplemented with 1X L-Glutamine, 10% heat-inactivated fetal bovine serum (FBS) and 100U penicillin/0.1 mg/mL streptomycin. All cells were grown at 5% CO_2_ and 37°C and regularly tested for mycoplasma. Transient transfection was performed using Lipofectamine RNAi max (Invitrogen) following manufacturer’s instructions. NR2F1-knockout cell lines were generated using CRISPR-Cas9–targeted genome editing Briefly, two separate NR2F1 gRNAs (guide 2, 5′-GATCCGCAGGACGACGTGGC-3′; and guide 4, 5′-GGCTGCCGTAGCGCGACGTG-3′) were cloned into pLentiCRISPRv2 (Addgene- 52961). An NT gRNA (5′-GTATTACTGATATTGGTGGG-3′) was used as a control. Lentiviral vectors were produced in HEK293T cells and viral supernatants were used to infect OMM1.3 QR^+^ cells. Cells were then selected using 2 µg/ml puromycin (Sigma-Aldrich; P8833). NR2F1 knockout was confirmed by Western blot. For NR2F1- inducible UM004 cell line, NR2F1 WT cDNA was cloned into pENTR/D-TOPO and then pLentiHygro3/TO/V-5-DEST. HEK293 cells were transfected with plasmid and generated lentivirus were used to infect UM004 cells. Expression of NR2F1 was induced using 0.1mg/ml and 1mg/ml of doxycycline for 48 h. For YAP1 overexpression, pLentipuro/TO/V5-DEST was used and overexpression of YAP1 was induced using 1mg/ml doxycycline in cells. YM-254890 was purchased from Focus Biomolecules. C26 (NR2F1 agonist) compound was purchased from Molport and required working stock concentration were prepared using 1mM primary stock of C26 prepared in DMSO to treat OMM1.3 cells.

### 2. Quantitative real-time PCR

RNA was isolated using RNeasy Plus Mini Kit (Qiagen), and cDNA was prepared using Superscript First Strand Synthesis System for RT-PCR Kit (Bio-Rad) following manufacturers’ instructions. Quantitative real-time PCR was performed using manufacturers’ instructions for SSO advanced SYBR green kit (Bio-Rad) or PowerUP SYBR green (Thermofisher scientific) on ABI-qPCR system using equal amount of template (50–100 ng cDNA) per reaction. Obtained values were normalized to housekeeping gene (Tubulin/RPL30/GAPDH) and are presented as fold differences over control or relative mRNA expression using the ΔΔCt/Livak method. Primer sequences are listed in supplementary table (Table 4A). For each experiment minimum sample size is (N=3) and p-values were calculated using student’s t-test.

### 3. Cell sorting

Livers sections were minced and digested with collagenase supplemented with DNase1 enzyme at 37^0^ C for 30 minutes followed by RBC lysis and percoll gradient was carried out isolated UM cells. Collected cells were resuspended in FACS sorting buffer and sent at Albert Einstein FACS core facility for cell sorting (based upon H2B-GFP^+/-^/td-tomato^+^ signal) using appropriate controls.

### 4. Western blot analysis

Cells were lysed in ice cold RIPA buffer (Thermofisher) supplemented with protease inhibitor cocktail Protease Inhibitor Cocktail (PIC) from Invitrogen. Each sample protein concentration was calculated using BCA protein assay kit and a standard BSA curve (Pierce). Equal amount of protein from each cell lysate and lysates were boiled in 4X Laemmli sample buffer for 5 minutes at 95°C followed by snap chill on ice. Samples were run on Bio-rad precasted gel in 1X running buffer (Bio-Rad) and transferred to PVDF membranes in 1X transfer buffer (Bio-rad) according to manufacturer’s protocol. Membranes were blocked in 5% non-fat milk or BSA in TBST (Tris-buffered saline supplemented with 0.1% Tween-20) buffer. Membranes were incubated with respected primary antibodies as indicated in each figure at 4°C with overnight shaking. Membranes were washed in TBST buffer and incubated with HRP-conjugated secondary antibodies at room temperature for 1 hour. Western blots were developed using pierce supersignal ECL or supersignal West Femto kit on iBright1500 alpha imager (Invitrogen). The antibodies used are listed in supplementary table (Table S4B). For densitometric analysis (N=3) independent blots were analyzed using ImageJ program and p-values were calculated using t-test.

### 5. Co-immunoprecipitation assay

Sub-confluent cultures of cells were harvested and lysed in ice cold immune-precipitation (IP) lysis buffer (Thermofisher) supplemented with PIC. Protein concentration was estimated using BCA kit as described above and total protein (0.7mg–2 mg) from each sample was immunoprecipitated with respected antibody using amounts as suggested in the manufacturer’s datasheets. Briefly, immunoprecipitated proteins were collected using 40 μl of preblocked protein A/G agarose beads and each immune precipitate was washed thrice with ice cold IP lysis buffer supplemented with PIC and finally beads were eluted in 50 μl of 2X loading dye. Samples was fractionated and immunodetection was done with anti-YAP1, TEAD1, NR2F1 and LATS1 antibodies at a dilution of 1:1000 (SI Table 4). For each immunoprecipitation IgG was used as internal control. Western blots were developed using pierce supersignal ECL or supersignal West Femto kit on iBright1500 alpha imager. For TEAD1and NR2F1 immunodetection membranes were incubated for 1 hour at room temperature with TidyBlot western blot detection reagent (Bio-rad) and for YAP1 HRP-conjugated secondary antibody was used.

### 6. Three-dimensional (3D) cell morphology studies

3D cell culture assays were performed in either 08-well chambers (Falcon) or in 06 well plates by coating with 80ul or 400-500 µl/well of Matrigel™ Matrix Growth Factor Reduced (R&D, Cultrex) respectively. Cells were suspended in 5% DMEM-F12 supplemented with 2% Matrigel™, plated at a density of 500- 1000/well for 08-well chamber and 0.5 x 10^4^ to 1 x 10^5^ in 06-well plate for different UM cell lines and incubated at 37°C for 5–14 days. A fresh layer of 5% DMEM-F12 supplemented with 2% Matrigel™ was added following 2 days of incubation, and the medium was replaced every 48 h with fresh DMEM-F12 medium. Colonies formed were monitored daily, and images were captured at either 5, 7 or 14 days using inverted phase contrast microscope (Nikon TS100). For label retention, colonies were counted (epifluorescence microscope) for GFP and td-tomato signal and ratio of GFP^+^/Td-tomato positive colonies were calculated. For colony counting, > 8-cell or <8- cell colonies were counted under the microscope and % event was calculated for each dataset data is plotted using graph pad Prism software with appropriate statistical test. For calculation of % event of colonies in 3D, minimum (N=3) independent samples were analyzed for calculating statistical significance except for fig. 6J where N=2.

### 7. Immunofluorescence

OMM1.3 and UM004 cells were cultured in 8-well chambers (Falcon) and 1000 cells/well were seeded in quadruplicate in 3D Matrigel (R&D, cultrex) and cells were fixed with 2% paraformaldehyde/PBS for 20 minutes at room temperature. The rest of the steps were completed as described by Khalil et al^55^ with minor modifications.

For immunofluorescence on paraffin-embedded tissue sections (FFPE), slides were incubated in xylene (05 minutes twice) followed by graded ethanol rehydration (3 minutes each), and finally washed with water for 5 min twice. Antigen retrieval for was performed in 10 mM antigen retrieval buffer (pH 6.0) from Invitrogen for 45 minutes using a steamer. Tissues were permeabilized in 0.3% TritonX-100-PBS for 05 minutes followed by three washes with 1X PBS and blocking in 0.1% TritonX-100-PBS supplemented with 5% normal goat serum and 3% bovine serum albumin (BSA, Fisher Bioreagents) at room temperature. Primary antibodies (listed in key table S4B) were diluted in blocking buffer (at recommended dilutions) and incubated overnight at 4°C followed by washing with PBS supplemented with 0.1% of TritonX-100- PBS three times. Cells were incubated with alexa fluor conjugated secondary antibodies at room temperature for 1-2 hour in the dark. Fixed slides were washed three times with 1X PBS and finally rinsed with distilled water. Slides were mounted with ProLong Gold Antifade reagent with DAPI. Confocal microscopy was performed using Leica SPE confocal microscope and images were analyzed to calculate MFI using ImageJ or Qupath software. For IF, minimum (N=3) independent field view or samples were included for analysis.

### 8. RNA Sequencing

Cells from OMM1.3 control and NR2F1_knockout cells cultured on 3D Matrigel for 07 days were extracted from Matrigel using cell dissociation buffer with continuous rocking at 4^0^ for 2-6 h. Cells were collected in ice cold 1X PBS and centrifuged at 300 g and cell pellets was resuspended in RLT lysis buffer. RNA was isolated using Qiagen RNeasy plus micro kit following manufacturer’s instructions. RNA samples QC, library preparation and RNA sequencing were performed at institutional genomics core facility. Qiaseq directional RNAseq kit (334387) with HMR rRNA removal (334386) and Qiaseq UDI Y-Adapter Kit was used to prepare RNA-seq libraries as per the manufacturer’s protocol. The samples were sequenced in a 2 x 75 bp paired-end configuration using the Illumina Nextseq 500 platform. Raw sequence data generated from Illumina NextSeq 500 were converted to FASTQ files and Quality control analysis was performed on fastq files using FastQC (https://www.bioinformatics.babraham.ac.uk/projects/fastqc/). Reads were aligned to the GRCh38 reference genomes using Star aligner ^81^ and GENCODE ^82^ gene and transcript annotations. For the *in vivo* dataset, RNA from sorted cells was isolated using NEBNext^®^ Single Cell/Low Input RNA Library Prep Kit. The samples were sequenced in a 2 x 150 bp paired-end configuration and generated reads were aligned to GRCm38, and XenofilteR ^83^ was used to remove any reads of mouse origin. RSEM ^84^ was used to quantify gene and transcript expression levels. Principal component analysis (PCA) and gene differential expression analysis was performed using DESeq2 ^85^. A paired-sample model was used for the *in vivo* dataset to eliminate any mouse-specific effects. PCA plots were rendered via the ggplot2 package (https://ggplot2.tidyverse.org). Heatmaps of normalized gene expression data were generated using the pheatmap package (https://cran.r-project.org/package=pheatmap). Data were analyzed in R (https://www.R-project.org/). Enrichr was used to determine significantly altered pathways ^86^. For RNA-seq, data analysis was carried out using (N=3) independent samples.

### 9. Cut & Run data analysis and data visualization

OMM1.3 control and NR2F1 knockout cells cultured in 3D were isolated using cell dissociation buffer as explained above in section 8. For each set, 30000 cells were collected and pooled as described in the Epicypher manual. The amount of Metstat panel was adjusted according to the number of cells used. Immunoprecipitated DNA was eluted in elution buffer and quantified on Qubit and used for library preparation and sequenced on Illumina NextSeq 2000 platform. As recommended, (2 x 50 cycles) paired end sequencing was performed. Raw data generated from Cut&Run paired-end sequencing reads were trimmed with trim galore (v0.6.7; https://github.com/FelixKrueger/TrimGalore) and then mapped to the mouse genome (mm10) using the Bowtie2 (v2.4.5; option -X 2000 –no-mixed –no-discordant). Read pairs with a mapping score >20 was kept for further analysis and duplicated reads due to PCR amplification were removed by SAM tools (v1.9). MACS2 (v 2.2.7.1) were used to call H3K4me3 and H3K27ac peaks in the paired-end mode and default options but for H3K27me3 enriched regions a sliding window method was used. Peaks from control and knockout samples were merged to generate a list of union peaks. Reads from individual samples located at the merged peaks were computed and used for sample clustering and differential modification analysis using the package EdgeR. To compare all three types of H3 modifications together, we counted the numbers of reads in each sample located to the +/- 2000 bp of transcription start sites of all the RefSeq genes and used them to perform principal component analysis. The software deep Tools (v3.5.1) was used for generating heatmaps of read densities centered on peaks, with data normalized to sequencing depths of 10 million. Read coverage was computed with the IGV tools and used for track visualization at IGV (only one replicate is shown). For peak analysis and pathways data coming from (N=3) samples was included in the study.

### 10. RNA interference/ Overexpression assays

OMM1.3, UM004 and 92.1 cells were transfected with gene-specific siRNAs with recommended concentration in 6-well using Lipofectamine RNAi Max reagent (Invitrogen) according to manufacturer’s instructions (Invitrogen). Briefly, cells were treated in 2D or 3D with either control, NR2F1 or YAP1 siRNA for 48h and for 05 days in 3D (new siRNA is replenished every 24h) and later cell lysates were harvested to confirm knockdown at either protein or transcript level using gene specific primers or antibodies. For YAP1 overexpression, 1μg of plasmid was used using similar protocol as described above except that after 48h of transfection, cells were treated with doxycycline (1ug/ml) to induce YAP1 overexpression. For all RNAi/ OE experiments minimum sample size is (N=3).

### 11. Chromatin immunoprecipitation (ChIP)

OMM1.3 and UM004 control and NR2F1 knockout and NR2F1 overexpressed cells were cross-linked with 1% PFA, washed, and lysed according to manufacturer’s protocol (EZ-ChIP; Millipore 17–371) with minor modifications. Briefly 1 x 10^6^ millions of either OMM1.3 or UM004 crosslinked cells were sonicated using Qsonica bioruptor (Diagenode) with power setting of 50 and 30 cycles of 30 s on/off. For each IP, 25 μg of chromatin was used for pull-down using antibodies specific to NR2F1, H3K4me, H3K27ac, H3K9me3 and corresponding isotype control. DNA was extracted from each immunoprecipitated and input samples and equal amount of DNA was used as template for qPCR using primers specific to promoter fragments of the YAP1 and control primers. Control primers (EZ-ChIP 22–004; Millipore) for the human GAPDH and was used as an internal control. Primer pairs were designed using NCBI primer designing tool and sequences of primers are listed in supplementary table (Table 4A). For enrichment of NR2F1 and H3K4me3 and H3K27ac marks (N=4) and (N=3) for H3K9me3 mark.

### 12. *In vivo* studies

BALB/nu female mice (7 weeks old) were purchased from the Charles River Laboratories and maintained under specific pathogen-free (SPF) conditions. All experiments were carried out according to IACUC guidelines. 2 × 10^6^ of OMM1.3-QR and UM004 control and NR2F1 knockout or overexpressed cells were resuspended in 30 μl of PBS and using aseptic condition, the spleen is accessed through a skin incision made by using a surgical blade, followed by access into the abdominal musculature by using surgical scissors near right lateral recumbency under gas anesthesia. Once the spleen is exposed, using a 27-gauge insulin syringe, cells were carefully injected into the spleen of mice respectively. Animals were monitored as per the IACUC guidelines and groups of mice were euthanized as per the experimental requirement. For metastatic burden and disseminations studies (N=6).

### 13. Human sample

Slides from Paraffin-embedded sections from clinically diagnosed UM patients with liver metastasis were obtained from Institute of Curie, Paris and Thomas Jefferson university, Philadelphia as per MTA guidelines. Samples received were deidentified and obtained with institutional review board approval. FFPE samples were stained for HMB45, NR2F1, YAP1 and Ki67. Nuclei were counterstained with DAPI. Several fields of view were assessed under each condition and to track UM cells (either solitary, < 8-cell cluster or > 8-cell cluster). Mean fluorescent intensity (MFI) was measured for each protein using QuPath software and HMB45 signal was used to track UM cells. Total 10 patient samples out to 20 were included in the study as some patient samples were not of good quality or failed during immunostaining step while others that did not show solitary or <8-cell cluster were also excluded from the study. For YAP1 quantification, only nuclear YAP1 and NR2F1 signal was included in quantification. Qupath software was used for quantification and only cells co-expressing HMB45 were included for analysis. For cumulative NRF1 and YAP1 in human patient sample size is (N=10) and several fields of views for each patient was used to plot the data.

### 14. Statistical analysis

To ensure reproducibility, *in vitro* experiments were repeated at least three times unless otherwise indicated. Data were expressed as means ± SDs from at three independent experiments. Depending on the experimental design, p-values were calculated using the unpaired Student’s *t*-test, multiple *t-test*, Mann– Whitney, or one-way ANOVA analysis as indicated. A p-value of < 0.05 was considered significant. Graph Pad Prism software was used to generate the graphs.

### 15. Data availability

Raw data generated from RNA-Seq and Cut&Run will be available on repository and other data generated in this study are available upon request from the corresponding author.

## Supporting information

Supplementary data (figure legeneds, figures, tables)

## Acknowledgements

We thank the Aguirre-Ghiso and Aplin labs as well as the Gutkind lab and Dr. Liliana Ossowski for helpful discussions. This work was supported by The National Institute of Health (NIH) /National Cancer Institute (NCI) (CA253977, CA109182, CA216248, CA218024, CA013330), DOD-MRP ME200152,

ME200270P2, the Samuel Waxman Cancer Research Foundation Tumor Dormancy Program and The Mark Foundation Aspire program and The Gurwin Foundation to J.A.A.G, who is also a Samuel Waxman Cancer Research Foundation Investigator. M.L.A. was recipient of a of a contract from Generalitat Valenciana (APOSTD2019/053). J.A.A.G is also supported by a grant from the Falk Foundation.

## Author’s contribution

R.K. planned, designed, performed experiments, analyzed data, conceptualized and wrote the manuscript. M.L.A planned and conducted experiments and analyzed data. T.J.P designed and analyzed RNA-seq data. V.C generated inducible cell lines and performed experiments. L.S contributed to human patient data quantification/analysis and also contributed to revising manuscript text. F.W.J generated YAP1 plasmids. J.L. F. T developed reagents and performed experiment. M.A.N wrote and edited the manuscript, provided scientific advice, and secured funding. T.S, M.T, R.S.S & L.D.K provided access to human samples (FFPE slides) used in this study. D.Z analyzed Cut & Run data. J.A.G conceived the project, designed the experiments along with A.E.A. J.A.G supervised the project, secured funding, and wrote the manuscript.

## Abbreviations

UM: Uveal melanoma
DCCs: Disseminated Cancer Cells
GPCR: G-protein coupled receptor
QR: Quiescence reporter.

