## Supplementary data (figure legeneds, figures, tables) for "Lineage commitment pathways epigenetically oppose oncogenic Gαq/11-YAP1 signaling in dormant disseminated uveal melanoma": Supplementary figure legends PDF_RK UM 01282025.pdf

[1]Department of Cell Biology, Montefiore Einstein Comprehensive Cancer Center, Albert Einstein College of Medicine, Bronx, NY, USA.
[2]Division of Hematology and Oncology, Department of Medicine, The Tisch Cancer Institute, Icahn School of Medicine at Mount Sinai, New York, NY.
[3] Instituto de Neurociencias (CSIC-UMH) and CIBERER, Alicante, Spain.
[4]Department of Pharmacology, Physiology, and Cancer Biology and the Sidney Kimmel Comprehensive Cancer Center, Thomas Jefferson University, Philadelphia, PA 19107, USA.
[5]School of Medical and Health Sciences, Edith Cowan University, Joondalup, WA 6027, Australia. Centre for Precision Health, Edith Cowan University, Joondalup, WA 6027, Australia
[6] Department of Medical Oncology, Thomas Jefferson University, Philadelphia, PA.
[7]Department of Translational Research, Institut Curie, PSL Research University, Paris, F-75005, France.
[8] Departments of Genetics, Neurology and Neuroscience, Albert Einstein College of Medicine, Bronx, NY, USA.
[9]Cancer Dormancy Institute, Montefiore Einstein Comprehensive Cancer Center, Albert Einstein College of Medicine, Bronx, NY, USA.
[10]Gruss-Lipper Biophotonics Center, Albert Einstein College of Medicine, Bronx, NY, USA.
[11]Ruth L. and David S. Gottesman Institute for Stem Cell Research and Regenerative Medicine, Albert Einstein College of Medicine, Bronx, NY, USA
[12]Institute for Aging Research, Albert Einstein College of Medicine, Bronx, NY, USA

[#] Present affiliation

Supplementary figure legends:

**Fig. S1.** A) Percentage of label retention (H2B-GFP) of OMM1.3 QR cells treated with doxycycline at day 0, day-7 and day-14 respectively. B) FACS plots representing OMM1.3 continuous cultured cells as control cells for gating td-tomato$^{+}$, GFP$^{+}$ (QR$^{+}$) and GFP$^{-}$(QR$^{-}$) population in presence and absence of doxycycline. C) FACS plots showing QR$^{+}$ and QR$^{-}$ UM cells population disseminated to the liver at different time points as indicated in the figure. These plots were used to calculate the percentage of disseminated UM cells as shown in Fig. 1C and for performing RNA-seq.

**Fig. S2.** A) Scatter plot showing the number of expressed genes versus the number of counts in the RNA-Seq data for QR$^{+}$ and QR$^{-}$ sorted sample from mice that passed aligned read cutoffs. The proliferative samples had approximately 5,000 more genes with expression and 12 million more transcripts than the quiescent samples. B) Pie chart showing distribution of differentially expressed genes in UM dormant DCCs (QR$^{+}$/QR$^{-}$) as analyzed from RNA-seq DESeq2 data. C) Expression of cell cycle related genes in terms of log2 fold change (p-adj.value < 0.05) in dormant UM DCCs population. D&E) GO terms predicted to be downregulated and upregulated by Enrichr KG in dormant OMM1.3-QR$^{+}$ UM DCCs using top 100 downregulated and upregulated genes in dormant UM DCCs population.

**Fig. S3.** A&B) Representative IF images and quantification graphs showing expression of NR2F1 (red) and Ki67 (red) in < and > 8-cell clusters 3 months post in UM004 and OMM1.3 (Ki67; grey) cells disseminated to the liver respectively. HMB45 was used to track UM cells. p-values calculated using Mann-Whitney test. C&D) IF images for Ki67 expression in < and > 8-cell clusters formed by OMM1.3 cells (magenta) at day 7 or in human metastatic UM patient samples (grey). Right panel for both images shows quantification of Ki67. p-values were calculates using Mann-Whitney test. E) Representative IF images co-stained for NR2F1 (green) and Ki67 (magenta) in < and > 8-cell clusters formed by OMM1.3 (upper panel) and UM004 cells (lower panel) respectively. Right panel shows quantification of nuclear NR2F1 and Ki67 M.F.I (A.U). F) Graphs showing nuclear NR2F1 MFI (A.U) in individual UM patient sample in < and > 8-cell clusters (N=10) quantified using Qupath software and p-value is calculated using Mann-Whitney test.

**Fig. S4**. A) Representative IF images co-stained for NR2F1(green) and YAP1 (red) in < and > 8-cell clusters formed by UM004 cells in 3D matrigel and quantification of total NR2F1 and YAP1 intensity per cell (A.U) in < and > 8-cell clusters is shown. B) Western blots showing basal expression of YAP1 in UM004, OMM1.3 and 92.1 UM cell line. GAPDH is used as loading control. C) mRNA expression of indicated genes treated with either control or NR2F1-targeting siRNA in OMM1.3 cells grown in 3D. p-values are calculated using student's t-test.

**Fig. S5**. A) mRNA expression of indicated genes treated with control or YAP1- siRNA in UM004 cells grown in 3D matrigel. p-values calculated by student's t-test. B) Western blot for YAP1 expression in OMM1.3 cell transiently transfected with control and YAP1-targeting siRNA. GAPDH was used as a loading control. C) Representative photographs of colonies formed by UM004 cells in 3D Matrigel at day 7 in control (No dox), NR2F1 OE (continuous dox treatment every 24h) and cell washed out after 48 h doxycycline condition. scale: 200x magnification and marked areas are shown as zoom in view of colony morphology. Quantification of % event of < and > 8-cell clusters formed under different conditions as shown in panel. p-values calculated using one-way anova. D) Graphs showing nuclear YAP1 MFI (A.U) in individual UM patient sample in < and > 8-cell clusters (N=10) as quantified by Qupath software and p-value is calculated using Mann-Whitney test.

**Fig. S6.** A) Expression of NR2F2 in OMM1.3 control and NR2F1 KO (KO1 and KO2) cell lysate. GAPDH was used as a loading control. B) Co-immunoprecipitation assay using lysates from OMM1.3 control and NR2F1 KO cells immunoprecipitated with anti-YAP1 antibody followed by immunodetection with anti-LATS1 antibody. IgG was used as isotype control. C) Representative image of colonies formed by OMM1.3

control and NR2F1 KO cells treated with either control or YAP1-siRNA at day 7. D) Representative images of liver surface imaging showing solitary OMM1.3-$QR^+$ DCCs in both control and NR2F1 knockout animals (N=6) and percentage of disseminated UM cells into the liver 36 h post intrasplenic injection.

**Fig. S7.** A) Heatmap showing log2-transformed normalized expression values of fifty significantly upregulated and downregulated genes in OMM1.3 control and NR2F1 KO RNA-seq samples based on unbiased clustering and p-adj values. B&C) GNAO1 mRNA expression in OMM1.3 control and NR2F1 knockout cells and UM004 WT and NR2F1 overexpressing cells grown in 3D Matrigel for 7 days. p-values calculated using student's t-test. D&E) Heatmaps generated from CUT&RUN data using H3K27me3 and H3K27ac antibody showing enriched and downregulated peaks for each condition.

**Fig. S8.** A&B) GO terms upregulated and downregulated by NR2F1 based on accumulation of H3k27me3 (cut off used = - 0.4 log2FC) or H3K27ac marks (cut off used = -1 log2FC) on genes in NR2F1 KO OMM1.3 cells used for Cut & Run analysis.

**Fig. S9.** A) Quantification of NR2F1 and YAP1 MFI (A.U) in colonies formed by OMM1.3 cells treated with DMSO and YM-300 nM at day 14. p-values calculated using Mann-Whitney test. B) Graph showing quantification of NR2F1 positive cells (< and > 8-cell clusters) following treatment with compound YM at different concentrations as indicated in the figure. C) Western blot showing expression of NR2F1 in OMM1.3 cells treated with 50 and 500 nM of NR2F1 agonist compound 26 at indicated time points. GAPDH was used as a loading control. * Indicates non-specific band D) Representative images of colonies at day 7 formed by OMM1.3 control cells and treated continuously for 5 days with compound C26 (50 nM) in 3D Matrigel at day 7. Right panel shows quantification of colonies (< and > 8-cell clusters) formed by OMM1.3 cells treated with either DMSO or C26 treated at 50 nM concentration (N=3). p-values calculated by student's t-test.

**Table S1.** RNA-seq data generated from $QR^+$/$QR^-$ sorted mouse samples (n=3) were used to generate DEG list. This table provides list of genes upregulated and downregulated in UM dormant DCCs with Gene ID, Log2FC with corresponding p-values (Tab 1; In vivo RNA-seq). Tab 2 and Tab 3 list genes that are upregulated and downregulated in dormant (QR+) UM DCCs. Tab 4,5,6 and 7 represents GO and KEGG terms downregulated and upregulated in dormant (QR+) UM DCCs. Tab 8 depicts genes that were used as background genes during analysis of GO and KEGG term using Enrichr database.

**Table S2A and S2B.** RNA-seq data generated NR2F1 WT (basal) and NR2F1 KO OMM1.3 cells grown in 3D Matrigel (n=3) were used to generate DEG list. Tab1 in this table provides list of genes upregulated and downregulated in NR2F1 KO group listed with Gene ID, Log2FC and corresponding p-values (3D RNA-seq). Table S2A represents GO terms activated in basal NR2F1 condition and Table S2B represents GO terms downregulated under basal NR2F1 conditions.

**Table S3A, S3B and S3B.** Differential peak for H3K27ac and H3K27me3 marks is shown in NR2F1 KO OMM1.3 cells grown in 3D. Table S3A shown GO terms repressed by NR2F1 due to loss of H3K27me3 marks on genes in NR2F1 KO group. Table S3B show GO terms activated by NR2F1 due to loss of H3K27ac marks on genes in NR2F1 KO condition. Table S3C shows GO terms repressed by NR2F1 due to loss of H3K27ac mark on genes in NR2F1 KO condition.
