## Supplementary data (figure legeneds, figures, tables) for "Lineage commitment pathways epigenetically oppose oncogenic Gαq/11-YAP1 signaling in dormant disseminated uveal melanoma": Supplementary figures PDF_1282025.pdf

[5]School of Medical and Health Sciences, Edith Cowan University, Joondalup, WA 6027, Australia. Centre for Precision Health, Edith Cowan University, Joondalup, WA 6027, Australia

[6] Department of Medical Oncology, Thomas Jefferson University, Philadelphia, PA.

[7]Department of Translational Research, Institut Curie, PSL Research University, Paris, F-75005, France.

[8] Departments of Genetics, Neurology and Neuroscience, Albert Einstein College of Medicine, Bronx, NY, USA.

[9]Cancer Dormancy Institute, Montefiore Einstein Comprehensive Cancer Center, Albert Einstein College of Medicine, Bronx, NY, USA.

[10]Gruss-Lipper Biophotonics Center, Albert Einstein College of Medicine, Bronx, NY, USA.

[11]Ruth L. and David S. Gottesman Institute for Stem Cell Research and Regenerative Medicine, Albert Einstein College of Medicine, Bronx, NY, USA

[12]Institute for Aging Research, Albert Einstein College of Medicine, Bronx, NY, USA

[#] Present affiliation

Fig. S1

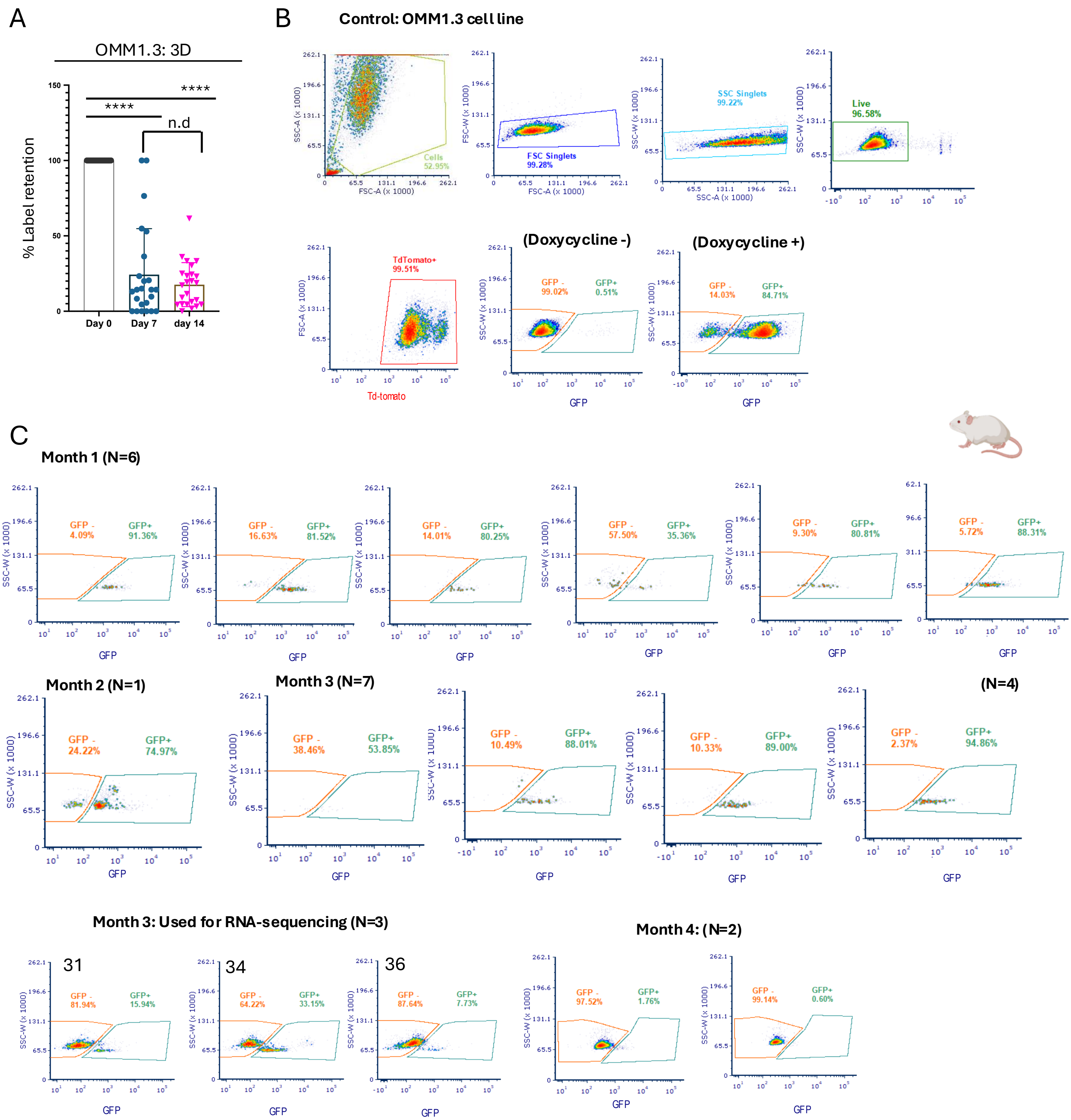

A
OMM1.3: 3D
% Label retention
****
****
n.d
Day 0
Day 7
day 14
B
Control: OMM1.3 cell line
Cells
52.95%
FSC Singlets
99.28%
SSC Singlets
99.22%
Live
96.58%
TdTomato+
99.51%
Td-tomato
(Doxycycline -)
GFP -
99.02%
GFP+
0.51%
(Doxycycline +)
GFP -
14.03%
GFP+
84.71%
C
Month 1 (N=6)
GFP - 4.09%
GFP+ 91.36%
GFP - 16.63%
GFP+ 81.52%
GFP - 14.01%
GFP+ 80.25%
GFP - 57.50%
GFP+ 35.36%
GFP - 9.30%
GFP+ 80.81%
GFP - 5.72%
GFP+ 88.31%
Month 2 (N=1)
GFP - 24.22%
GFP+ 74.97%
Month 3 (N=7)
GFP - 38.46%
GFP+ 53.85%
GFP - 10.49%
GFP+ 88.01%
GFP - 10.33%
GFP+ 89.00%
(N=4)
GFP - 2.37%
GFP+ 94.86%
Month 3: Used for RNA-sequencing (N=3)
31
GFP - 81.94%
GFP+ 15.94%
34
GFP - 64.22%
GFP+ 33.15%
36
GFP - 87.64%
GFP+ 7.73%
Month 4: (N=2)
GFP - 97.52%
GFP+ 1.76%
GFP - 99.14%
GFP+ 0.60%
GFP
SSC-W (x 1000)

Fig. S2

A

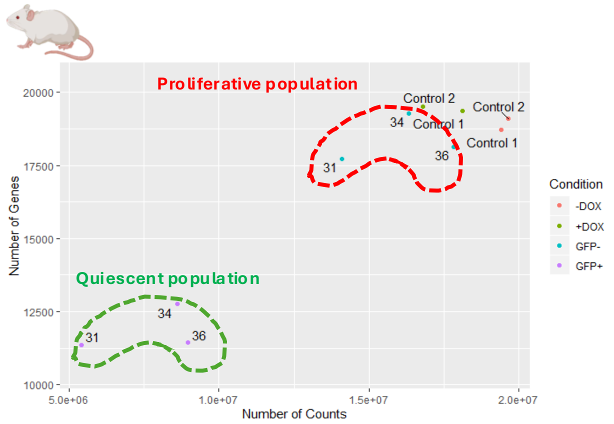


B

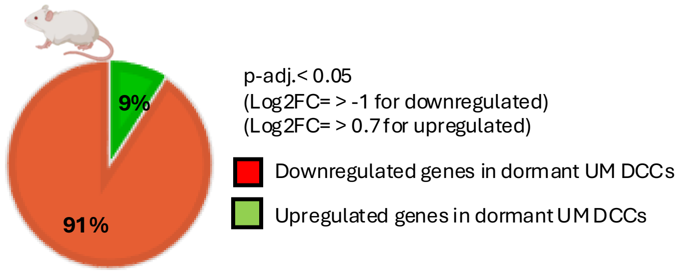


C

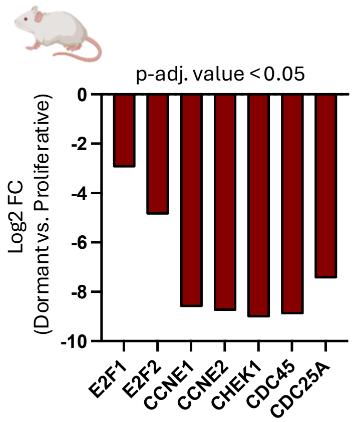


D

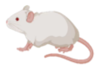

Pathways downregulated in Dormant UM population

| Term | Library | p-value |
|---|---|---|
| **positive regulation of neurogenesis (GO:0050769)** | GO_Biological_Process_2021 | 0.0004664 |
| **regulation of small GTPase mediated signal transduction (GO:0051056)** | GO_Biological_Process_2021 | 0.0007139 |
| **regulation of cell cycle G2/M phase transition (GO:1902749)** | GO_Biological_Process_2021 | 0.0008736 |
| **regulation of G2/M transition of mitotic cell cycle (GO:0010389)** | GO_Biological_Process_2021 | 0.0009152 |
| **DNA integrity checkpoint signaling (GO:0031570)** | GO_Biological_Process_2021 | 0.0009726 |

E

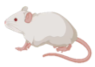

Pathways upregulated in Dormant UM population

| Term | Library | p-value |
|---|---|---|
| **positive regulation of glial cell differentiation (GO:0045687)** | GO_Biological_Process_2021 | 0.000093713 |
| **negative regulation of nucleic acid-templated transcription (GO:1903507)** | GO_Biological_Process_2021 | 0.0005304 |
| **positive regulation of astrocyte differentiation (GO:0048711)** | GO_Biological_Process_2021 | 0.0006796 |
| **negative regulation of DNA-binding transcription factor activity (GO:0043433)** | GO_Biological_Process_2021 | 0.0007607 |
| **regulation of astrocyte differentiation (GO:0048710)** | GO_Biological_Process_2021 | 0.001581 |

Fig. S3

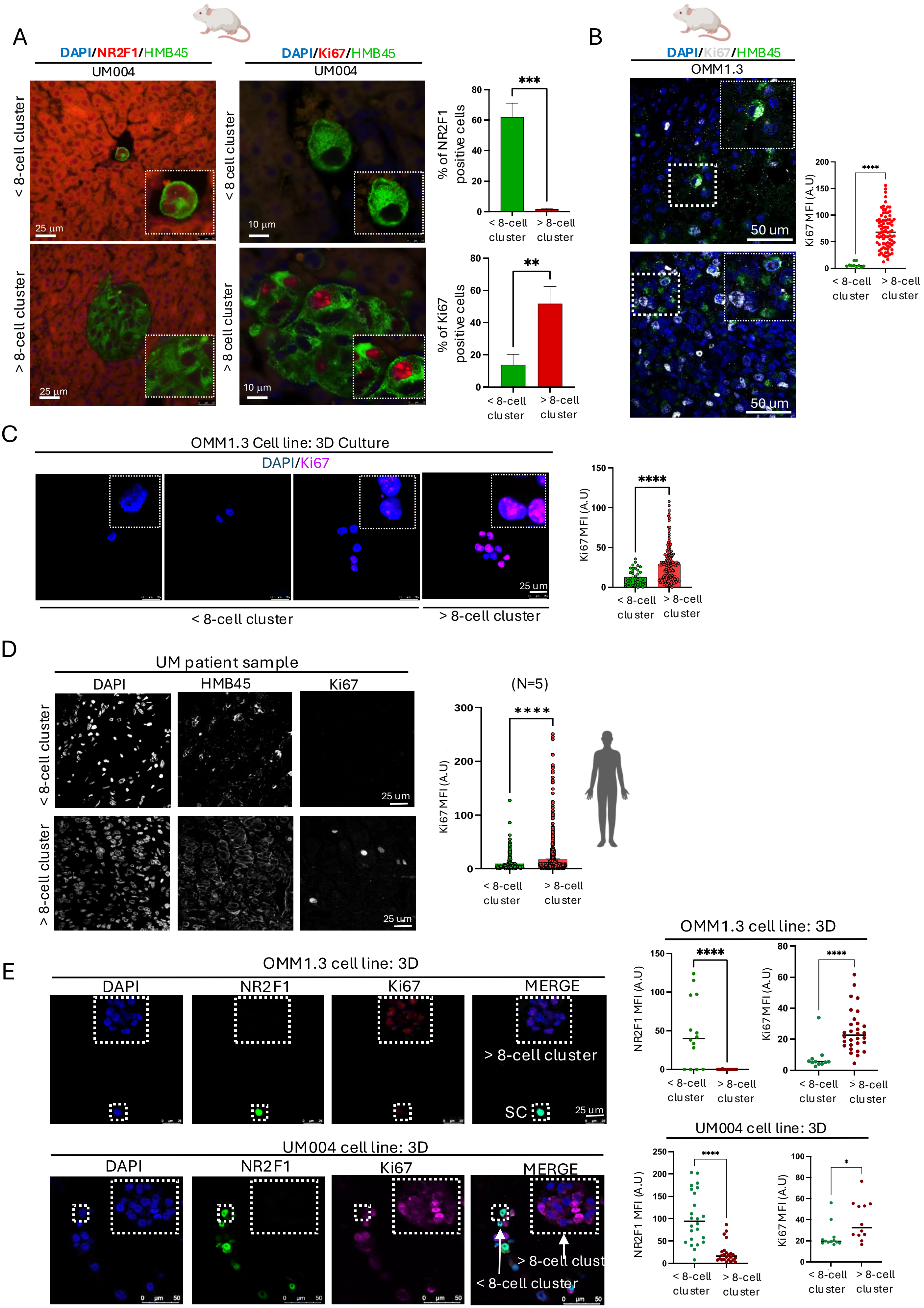
A
DAPI/NR2F1/HMB45
UM004
DAPI/Ki67/HMB45
UM004
< 8-cell cluster
> 8-cell cluster
25 µm
10 µm
% of NR2F1 positive cells
% of Ki67 positive cells
***
**
< 8-cell cluster
> 8-cell cluster
B
DAPI/Ki67/HMB45
OMM1.3
50 um
Ki67 MFI (A.U)
****
C
OMM1.3 Cell line: 3D Culture
DAPI/Ki67
25 um
< 8-cell cluster
> 8-cell cluster
D
UM patient sample
DAPI
HMB45
Ki67
(N=5)
Ki67 MFI (A.U)
E
OMM1.3 cell line: 3D
DAPI
NR2F1
Ki67
MERGE
> 8-cell cluster
SC
UM004 cell line: 3D
< 8-cell cluster
NR2F1 MFI (A.U)
*

Fig. S3F

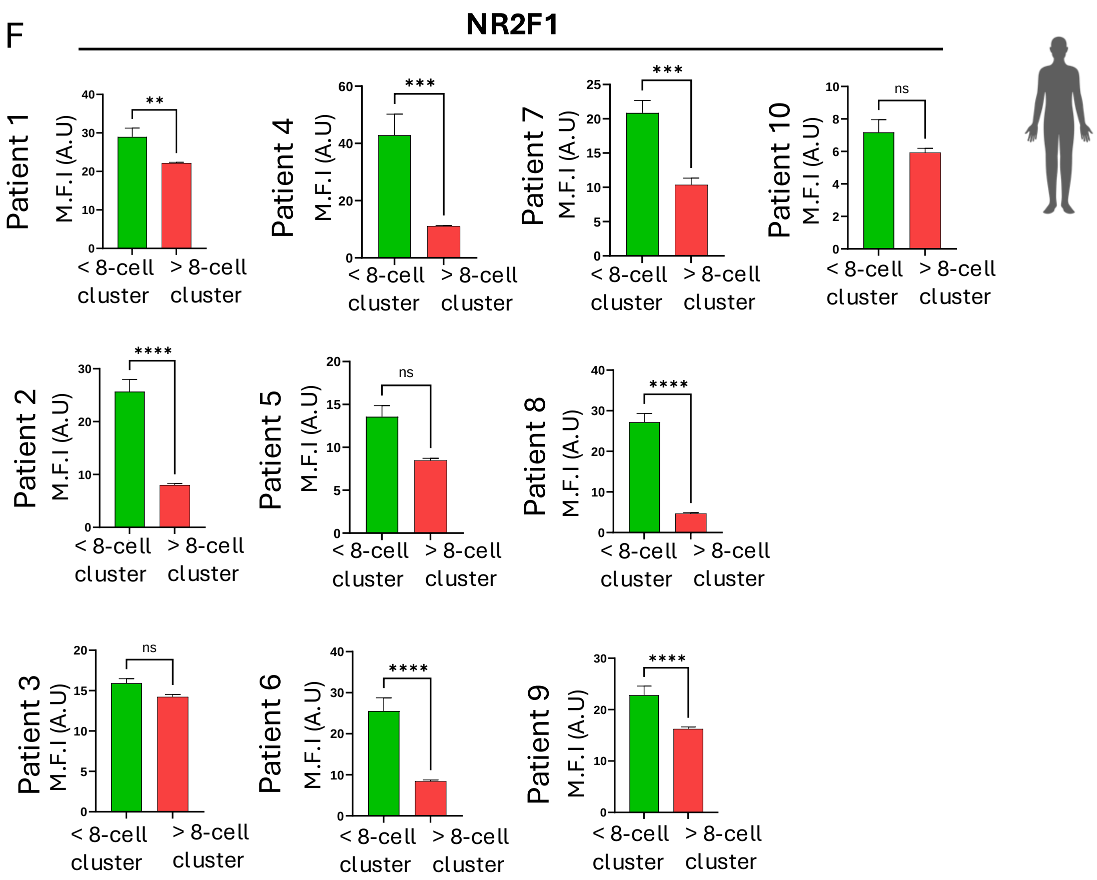
F
NR2F1
Patient 1
Patient 2
Patient 3
Patient 4
Patient 5
Patient 6
Patient 7
Patient 8
Patient 9
Patient 10
M.F.I (A.U)
< 8-cell cluster
> 8-cell cluster
**
***
****
ns

Fig. S4

A

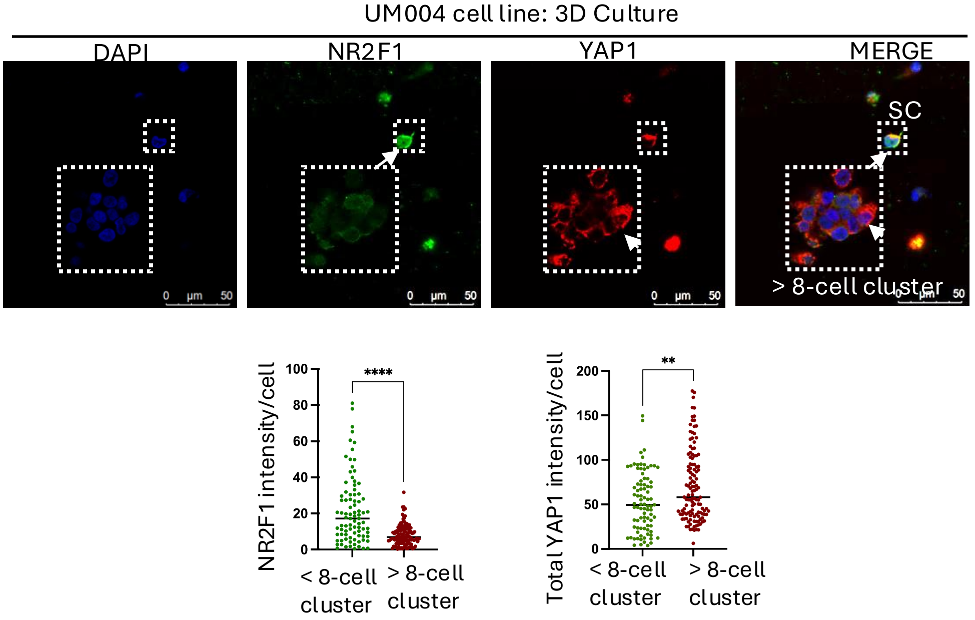
UM004 cell line: 3D Culture
DAPI
NR2F1
YAP1
MERGE
SC
> 8-cell cluster
0 µm 50
NR2F1 intensity/cell
****
< 8-cell cluster
> 8-cell cluster
Total YAP1 intensity/cell
**
< 8-cell cluster
> 8-cell cluster

B

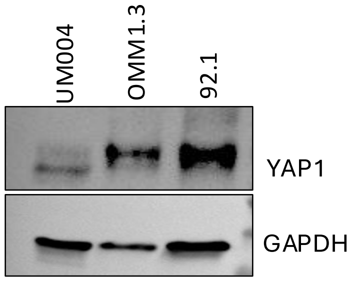
UM004
OMM1.3
92.1
YAP1
GAPDH

C

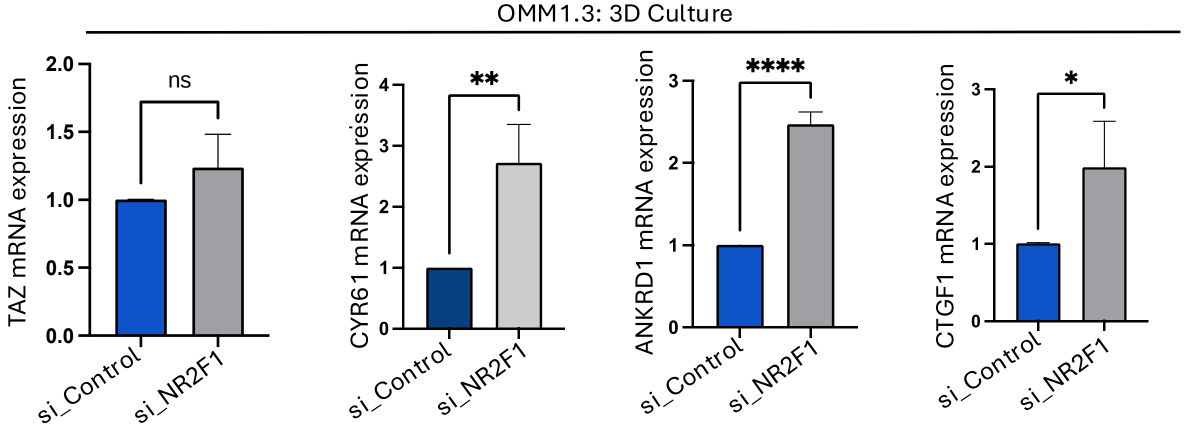
OMM1.3: 3D Culture
TAZ mRNA expression
ns
CYR61 mRNA expression
**
ANKRD1 mRNA expression
****
CTGF1 mRNA expression
*
si_Control
si_NR2F1

Fig. S5

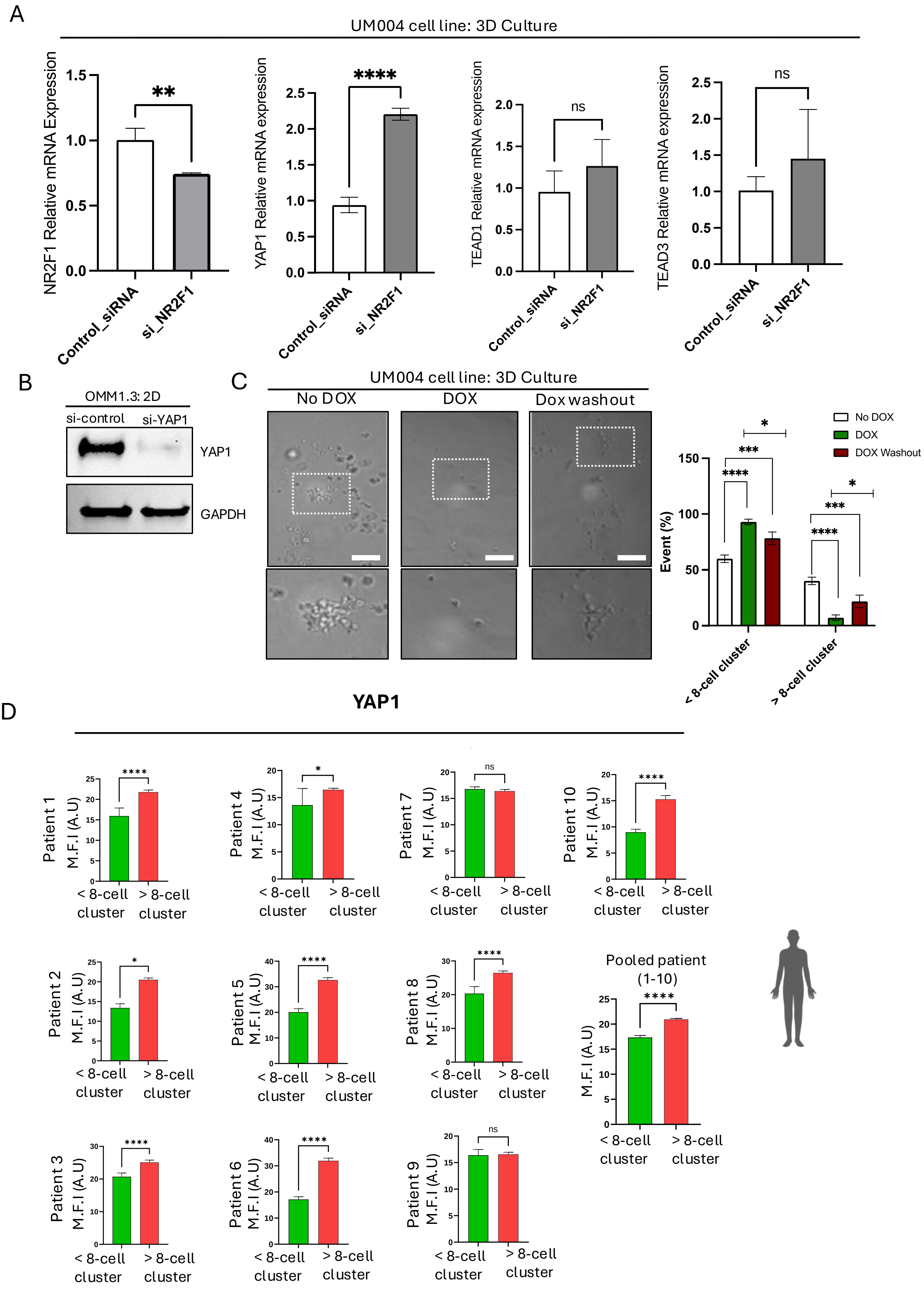
A
UM004 cell line: 3D Culture
NR2F1 Relative mRNA Expression
YAP1 Relative mRNA expression
TEAD1 Relative mRNA expression
TEAD3 Relative mRNA expression
**
****
ns
ns
Control_siRNA
si_NR2F1
B
OMM1.3: 2D
si-control
si-YAP1
YAP1
GAPDH
C
UM004 cell line: 3D Culture
No DOX
DOX
Dox washout
No DOX
DOX
DOX Washout
Event (%)
< 8-cell cluster
> 8-cell cluster
D
YAP1
Patient 1
Patient 2
Patient 3
Patient 4
Patient 5
Patient 6
Patient 7
Patient 8
Patient 9
Patient 10
Pooled patient (1-10)
M.F.I (A.U)
< 8-cell cluster
> 8-cell cluster

Fig: S6

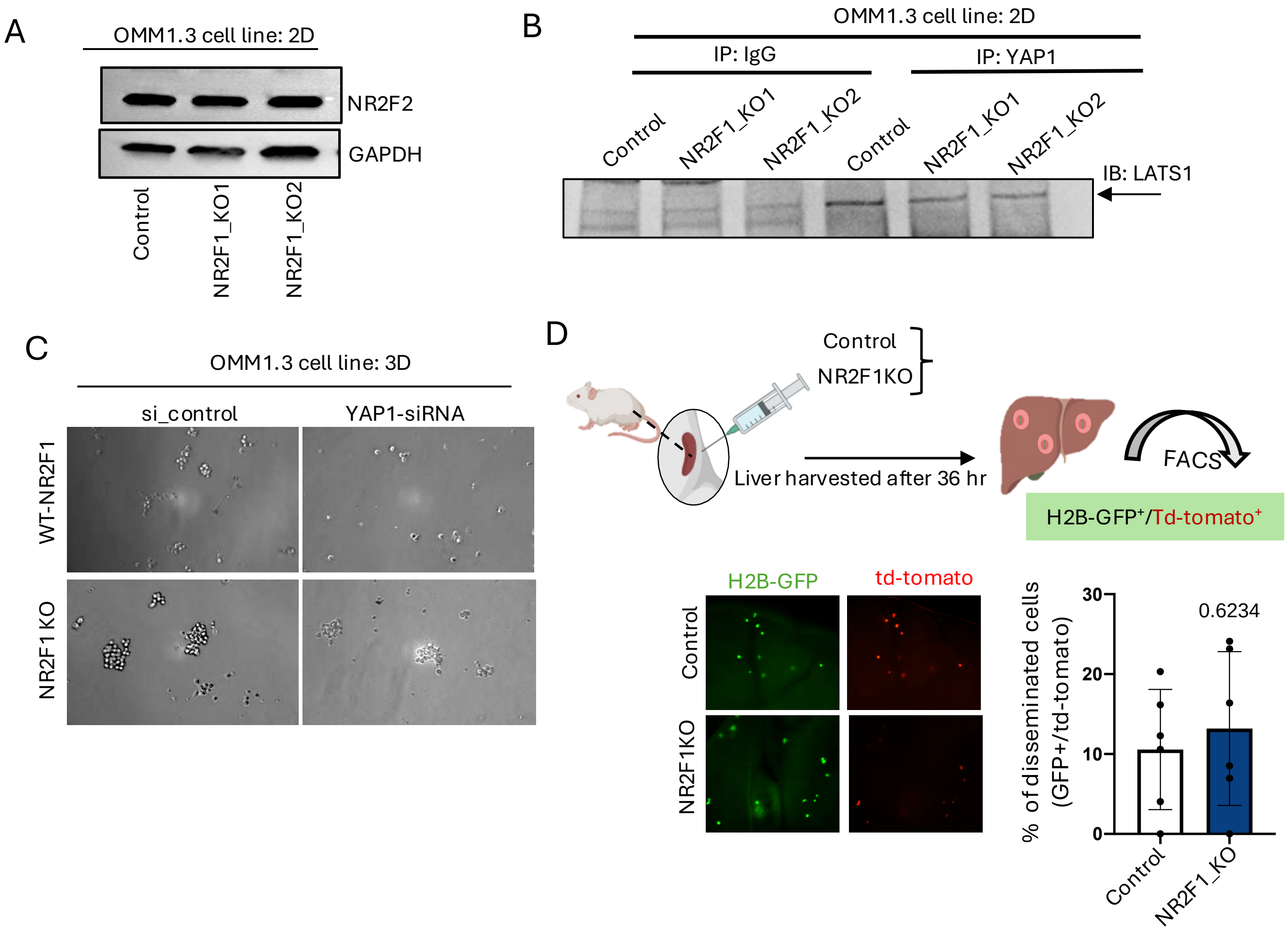
A
OMM1.3 cell line: 2D
NR2F2
GAPDH
Control
NR2F1_KO1
NR2F1_KO2
B
OMM1.3 cell line: 2D
IP: IgG
IP: YAP1
Control
NR2F1_KO1
NR2F1_KO2
Control
NR2F1_KO1
NR2F1_KO2
IB: LATS1
C
OMM1.3 cell line: 3D
si_control
YAP1-siRNA
WT-NR2F1
NR2F1 KO
D
Control
NR2F1KO
Liver harvested after 36 hr
FACS
H2B-GFP+/Td-tomato+
H2B-GFP
td-tomato
Control
NR2F1KO
% of disseminated cells (GFP+/td-tomato)
30
20
10
0
0.6234
Control
NR2F1_KO

Fig. S7

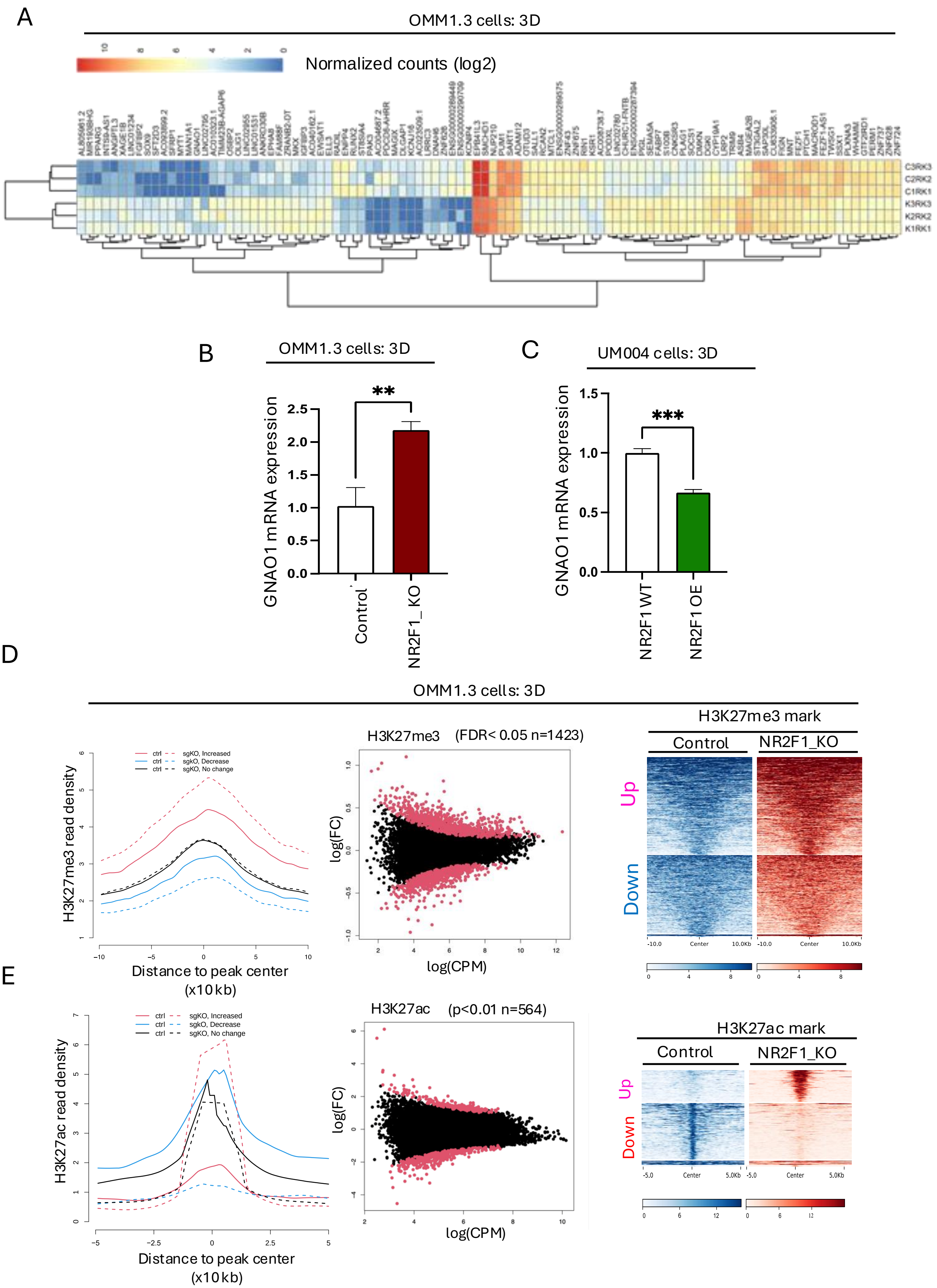
A
OMM1.3 cells: 3D
Normalized counts (log2)
C3RK3
C2RK2
C1RK1
K3RK3
K2RK2
K1RK1
B
OMM1.3 cells: 3D
GNAO1 mRNA expression
**
Control
NR2F1_KO
C
UM004 cells: 3D
GNAO1 mRNA expression
***
NR2F1 WT
NR2F1 OE
D
OMM1.3 cells: 3D
H3K27me3 read density
ctrl sgKO, Increased
ctrl sgKO, Decrease
ctrl sgKO, No change
Distance to peak center (x10kb)
H3K27me3 (FDR< 0.05 n=1423)
log(FC)
log(CPM)
H3K27me3 mark
Control
NR2F1_KO
Up
Down
E
H3K27ac read density
ctrl sgKO, Increased
ctrl sgKO, Decrease
ctrl sgKO, No change
Distance to peak center (x10kb)
H3K27ac (p<0.01 n=564)
log(FC)
log(CPM)
H3K27ac mark
Control
NR2F1_KO
Up
Down

Fig. S8

A

GO term activated by NR2F1 (H3K27me3 induced changes)

| GO Biological Process | p-value |
|---|---|
| Monoatomic Cation Transmembrane Transport (GO:0098655) | 0.000996635963537511 |
| Inorganic Cation Transmembrane Transport (GO:0098662) | 0.0010734180759699000 |
| Regulation Of Catecholamine Secretion (GO:0050433) | 0.0017604877931649200 |
| Anterograde Trans-Synaptic Signaling (GO:0098916) | 0.0021384881004009200 |
| Positive Regulation Of Lipid Kinase Activity (GO:0090218) | 0.0023960411238384900 |
| Chemical Synaptic Transmission (GO:0007268) | 0.003267357585211800 |
| Action Potential (GO:0001508) | 0.003738268800972130 |
| Ionotropic Glutamate Receptor Signaling Pathway (GO:0035235) | 0.004956243986150180 |
| Semaphorin-Plexin Signaling Pathway Involved In Neuron Projection Guidance (GO:1902285) | 0.004956243986150180 |
| Negative Regulation Of Cell Activation (GO:0050866) | 0.005823066766900990 |

B

GO term activated by NR2F1 (H3K27ac induced changes)

| GO biological Process | p-value | q-value |
|---|---|---|
| Nervous System Development (GO:0007399) | 2.71E-05 | 0.06496497 |
| Melanocyte Differentiation (GO:0030318) | 4.51E-05 | 0.06496497 |
| Positive Regulation Of Cellular Biosynthetic Process (GO:0031328) | 0.0001133 | 0.10887706 |
| Negative Regulation Of Sodium Ion Transmembrane Transport (GO:1902306) | 0.00031099 | 0.2161356 |
| Positive Regulation Of Cell Projection Organization (GO:0031346) | 0.00037484 | 0.2161356 |
| Positive Regulation Of Epithelial To Mesenchymal Transition (GO:0010718) | 0.00062524 | 0.25380218 |
| Negative Regulation Of Intracellular Signal Transduction (GO:1902532) | 0.00064291 | 0.25380218 |
| Melanosome Organization (GO:0032438) | 0.00070536 | 0.25380218 |
| Negative Regulation Of Cytoskeleton Organization (GO:0051494) | 0.00085205 | 0.25380218 |
| Peptidyl-Serine Phosphorylation (GO:0018105) | 0.00088034 | 0.25380218 |

Fig. S9

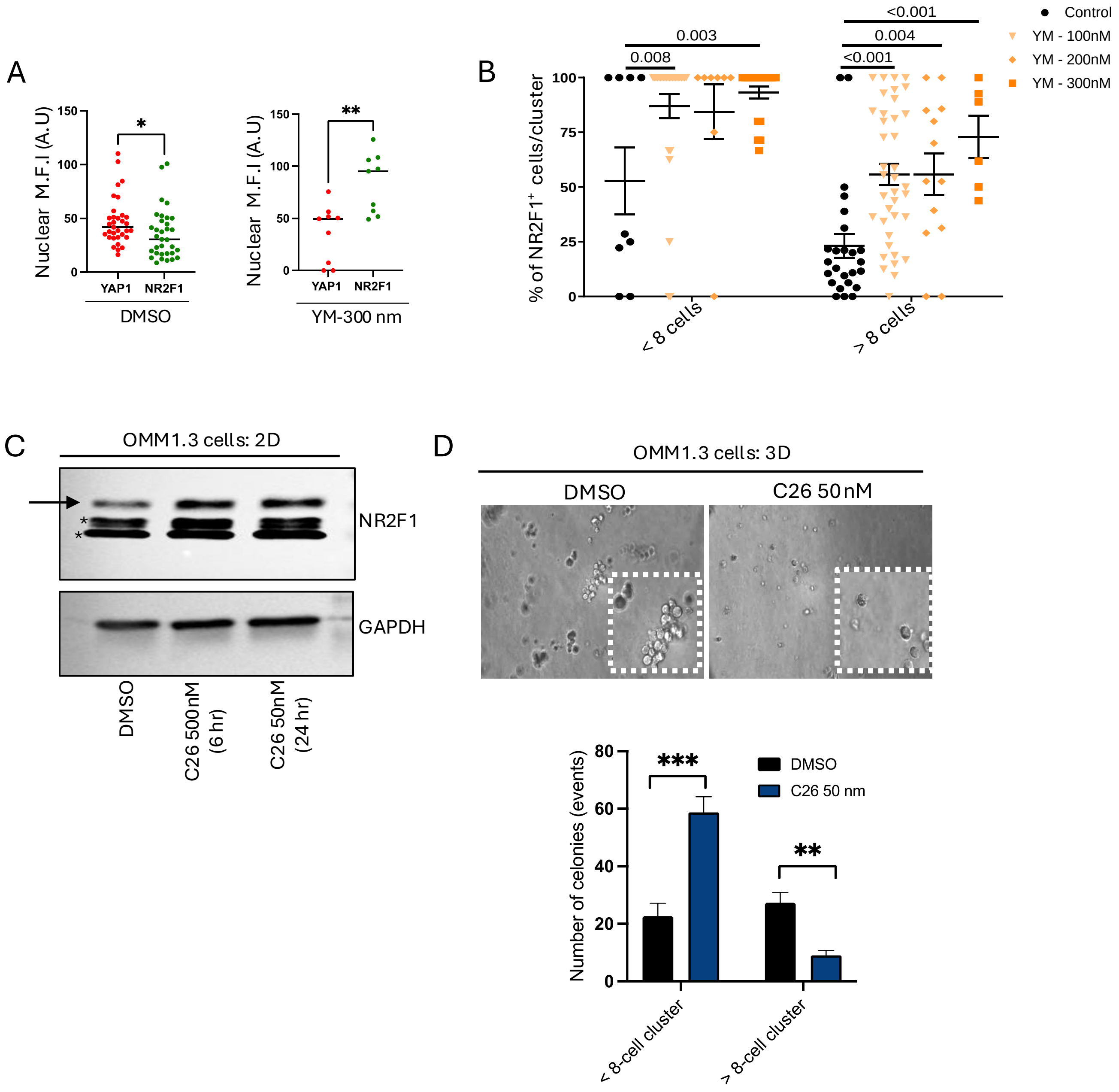
A
Nuclear M.F.I (A.U)
150
100
50
0
*
YAP1
NR2F1
DMSO
**
YM-300 nm
B
% of NR2F1+ cells/cluster
0.003
0.008
<0.001
0.004
<0.001
< 8 cells
> 8 cells
Control
YM - 100nM
YM - 200nM
YM - 300nM
C
OMM1.3 cells: 2D
NR2F1
GAPDH
DMSO
C26 500nM (6 hr)
C26 50nM (24 hr)
D
OMM1.3 cells: 3D
DMSO
C26 50nM
Number of celonies (events)
***
**
DMSO
C26 50 nm
< 8-cell cluster
> 8-cell cluster
